# Soil intake modifies the gut microbiota and alleviates ovalbumin-induced mice asthma inflammation

**DOI:** 10.1101/2023.05.31.543010

**Authors:** Mengjie Li, Na Li, Yangyang Dong, Honglin Zhang, Zhimao Bai, Rui Zhang, Zhongjie Fei, Wenyong Zhu, Pengfeng Xiao, Xiao Sun, Dongrui Zhou

## Abstract

**Background:** Low cleanliness living environment (LCLE) can increase gut microbial diversity and prevent allergic diseases, whereas gut microbial dysbiosis is closely related to the pathogenesis of asthma. Our previous studies suggested that soil in the LCLE is a key factor in shaping intestinal microbiota.

**Objective:** We aimed to explore if sterilized soil intake as prebiotics while being incubated with microbes in the air can attenuate mice asthma symptoms by modifying gut microbiota.

**Methods:** 16S rRNA gene sequencing was used to analyze the gut microbial composition, in combination with immune parameters measured in the lung and serum samples.

**Results:** 16S rRNA gene sequencing results showed significant differences in the fecal microbiota composition between the test and control mice, with a higher abundance of *Allobaculum*, *Alistipes,* and *Lachnospiraceae_UCG-001*, which produce short-chain fatty acids and are beneficial for health in the test mice. Soil intake downregulated the concentrations of IL-6, IL-4, IL-17F, TNF-α, and IL-22 in serum and increased the expression of IFN-γ, which regulated the Th1/Th2 balance in lung by polarizing the immune system toward Th1, strongly alleviating ovalbumin-induced asthma inflammation. The effect of sensitization on gut microbiota was greater than that of air microbes and age together, but weaker than that of soil.

**Conclusion:** Soil intake had a significant therapeutic effect on mouse asthma, possibly by promoting the growth of multiple beneficial bacteria. The results indicated that the development of soil-based prebiotic products might be used for allergic asthma management and our study provides further evidence for the hygiene hypothesis.

**Importance:** Exposure to a low cleanliness living environment (LCLE), of which soil is an important component, can shape the gut microbiota and support immune tolerance, preventing allergic diseases such as eczema and asthma. However, with the rapid progress of urbanization, it is impossible to return to farm-like living and we are becoming disconnected from the soil. Here, our study found that ingesting sterilized soil and living in an LCLE have the same protective effects on asthma inflammation. Ingestion of sterilized soil significantly altered the gut microbial composition and exerted significant therapeutic effects on asthmatic mice. However, edible sterilized soil possesses more advantages than LCLE exposure, such as the absence of pathogenic bacteria, safer, and convenience. The results indicate that the development of soil-based prebiotic products might be used for allergic asthma management and our study further supports the hygiene hypothesis.

**Notification:** The article is currently undergoing peer review in the World Allergy Organization Journal.

## Introduction

Asthma is a common chronic respiratory inflammatory disease with symptoms including shortness of breath, chest tightness, and coughing. It affects over 300 million people worldwide, a number that is expected to increase to 400 million by 2025(1).

Numerous epidemiological studies have shown that low cleanliness living environment (LCLE) exposure can protect against asthma and atopic diseases(2,3),(4). A previous study showed that mice housed in cages with soil, house dust, and decaying plant material had higher gut microbial diversity and lower serum immunoglobulin E (IgE) levels than specific pathogen-free (SPF) mice after administration of 2-4-dinitrofluorobenzene(5). Similarly, Ottman et al. found that mice housed in LCLE altered gut microbial composition and alleviated Th2-type allergic responses(6). Besides, LCLE exposure resulted in faster gut microbiota maturation in weaned piglets(7). Furthermore, keeping antibiotic-treated mice in LCLE accelerated gut microbial recovery(8). Recently, human studies showed that direct contact with the varied microbial community of soil– and plant-based material stabilized human gut microbes, triggering a beneficial immune response in allergic diseases(9).

Targeting the gut microbiota using probiotics(10,11), prebiotics(12), and fecal microbiota transplantation(13,14), positively affects asthma. Due to the diverse composition of the LCLE, it remains unclear which factors play a key role in its protective effect against asthma. Our previous study indicated that soil is a crucial factor in shaping intestinal microecology and that exposing mice to sterilized soil significantly changed the gut microbial composition(15). Furthermore, ingestion of sterilized soil, while inhaling microbes in the air, reduced serum IgE levels by increasing gut microbial diversity(16). However, it is unknown if sterilized soil intake as prebiotics can achieve a therapeutic effect against ovalbumin (OVA)-induced asthma by changing intestinal microbiota.

To explore the therapeutic effect of oral sterilized soil prebiotics on asthma and its effects on intestinal microorganisms and immune function, we established OVA-induced mouse asthma models in an SPF animal room. These mice were transferred to the general animal room, where they could be continuously inoculated with the microorganisms in the air. Mice in the test group were fed 5% sterilized soil. After 8-week soil intervention, 16S rRNA gene sequencing was used to analyze the intestinal microbiota composition. The effect of soil intake on the immune function of asthmatic mice was investigated using pathological section examinations, flow cytometry analyses, and Reverse transcription-quantitative polymerase chain reaction (RT-qPCR). As increasing evidence shows the bidirectional communication between the gut microbiota and systemic immunity along the gut–lung axis(17,18), we also further examined the differences in the intestinal microbiota composition between the steady-state and asthmatic mice.

## Methods

### Animals and asthma model

Adult male and female C_57_ Bl/6 mice with the same genetic background were purchased from B&K Universal (Shanghai, China) and housed in the SPF animal facility. All animal experiments were approved by the National Institute of Bioscience and the Advisory Committee on Animal Health Research of Southeast University (Approval number: 2019101522). Male mice (n = 23, 4-week-old) were selected from the offspring and co-housed until age 8 weeks. Then, they were randomly divided into P_Asthma (n = 15) and P_PBS (n = 8) groups. Detailed information regarding the mice and treatment is described (Table E1).

We employed the asthma modeling method proposed by Noora et al(6). with appropriate modifications (Fig. E1). P_Asthma group mice were exposed to the murine asthma model protocol, receiving two intraperitoneal injections of 50 μg OVA emulsified in 2.25 mg of alum in a total volume of 100 μL of PBS on days 0 and 14, followed by 5% aerosolized OVA challenge from days 21 to 27, for 40 min each time. P_PBS group mice received two intraperitoneal PBS injections and were challenged with PBS only. Mouse feces were collected on day 28. Eight mice were randomly selected from the P_Asthma group for soil intervention (Soil group, n = 8). The above operations were conducted in the SPF animal room. Then, all mice were transferred to the general animal room. Asthma and PBS group mice were fed a standard diet, while Soil group mice were fed a standard diet containing 5% sterilized soil. All conditions, including gender, age, house temperature, and humidity, were equal for the three groups. After 8-week soil intervention, Asthma and Soil group mice were challenged with 5% aerosolized OVA for 40 min while PBS group mice were challenged with PBS. Feces, lung, serum, and bronchoalveolar lavage fluid (BALF) samples were collected the next day for subsequent experiments. The detailed timeline of the experimental design is shown in Fig. E2.

### Soil and feed

The soil used in the experiment, which was ground into powder and filtered, was from the top 10 cm of farm ground, which is rich in vegetation with good ecology and unpolluted. Standard feed for mice (Xietong Pharmaceutical Bioengineering Company) was ground into powder using a flour mill. Sterilization was performed twice by autoclaving at 121 °C for 30 min. Sterilized soil (5 g) and standard feed (95 g) were weighed and then mixed with sterilized water on a super-clean bench. See Table S3 for a detailed soil composition analysis(16).

### DNA extraction and sequencing

DNA sequencing was performed at Shanghai Lingen Biotechnology Co., Ltd. Fecal microbial DNA was extracted using a fecal DNA kit (Omega Bio-tek, Norcross, GA, USA). The V4-V5 region of the bacterial 16S ribosomal RNA gene was amplified using PCR and 515F 5′-barcode-GTGCCAGCMGCCGCGG-3′ and 907R 5′-CCGTCAATTCMTTTRAGTTT-3′ primers. PCR products were quantified using the QuantiFluor™-ST Blue Fluorescence Quantification System (Promega). After Illumina PE250 library construction, sequencing of PCR product libraries was performed on the Illumina MiSeq platform (Illumina Inc, San Diego, CA, USA). The paired-end (PE) reads obtained from sequencing were spliced into a sequence according to the overlapping relationship between them, while the sequence quality was controlled and filtered. Using the QIIME software package (V1.9.0 http://qiime.org/scripts/assign_taxonomy.html), sequences with similarities ≥ 97% were clustered into an operational taxonomic unit (OTU) and analyzed taxonomically for species.

### Real-time quantitative PCR

Total RNA was extracted from mouse lung tissue using an RNA extraction kit according to the manufacturer’s instructions (BioMiGA, China). All materials required for the experiment were treated with DPEC water to remove RNA enzyme (BioMiGA, China). cDNA was obtained using reverse transcription PCR (Yugong Biolabs, China). The reaction system was prepared according to the kit’s requirements (Yugong Biolabs) and fluorescence quantitative analysis was performed using the ABI 7500 PCR system (Applied Biosystems). Conditions were as follows: 30 s at 95 L, 40 cycles of 10 s at 95 L, and 30 s at 60 L, followed by 15 s at 95 L, 60 s at 60 L, and 15 s at 95 L. The cytokine mRNA expression levels were calculated according to the 2^-ΔΔCT^ formula. Glyceraldehyde 3-phosphate dehydrogenase (GAPDH) was used as a control.

### Flow cytometry

The LEGENDplex™ MU Th Panel kit (Biolegend) was used to detect serum cytokine concentrations using flow cytometry (FACS Calibur). Quantitative analyte analysis was performed using the LEGENDplex v8.0 software.

### Data availability

The raw data of 16S rRNA gene sequences were deposited in the NCBI Sequence Read Archive (SRA) database (project number, PRJNA877056).

### Ethics approval

All animal experiments were performed in strict accordance with the guidelines of the Ethical Committee on Animal Research of Southeast University and this study obtained ethics approval from the National Institute of Bioscience and the Advisory Committee on Animal Health Research of Southeast University (Approval number: 2019101522).

## Results

### Soil intake shaped the gut microbial composition and promoted beneficial bacteria growth

Mice were transferred to the general animal room, where they were incubated with the air microorganisms (Fig. 1A). Soil group mice were simultaneously fed 5% sterilized soil. After the 8-week soil treatment, feces were collected for 16S rRNA gene sequencing.

**Figure 1.**
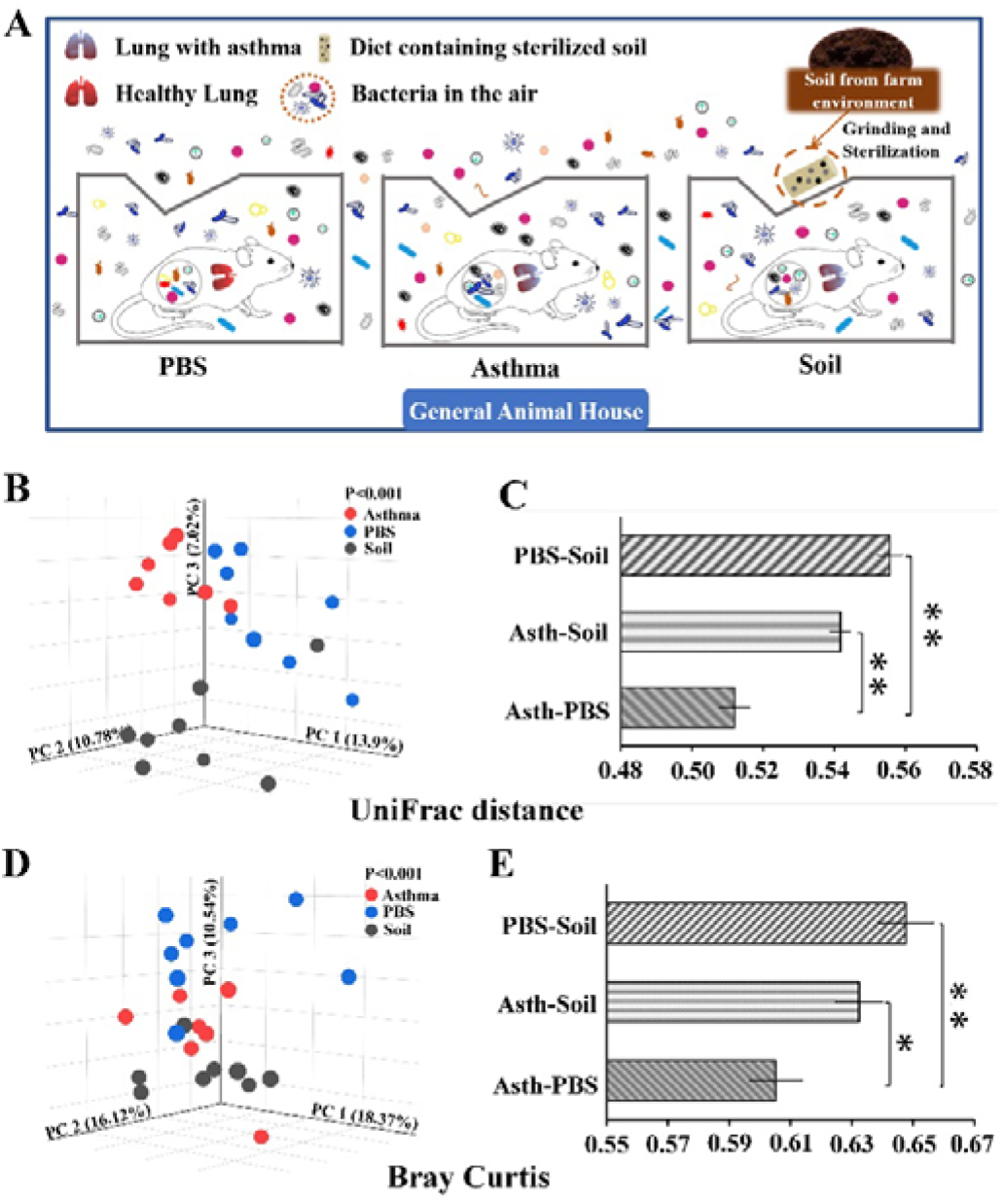
Schematic diagram of experimental treatments, and gut microbial beta diversity. A. Soil treatment mode for asthmatic mice: after completing the asthma model in the SPF animal room, all three groups of mice were transferred to the general animal room with microbes in the air. 5% sterilized soil was added to the feed of the Soil group. B. PCoA of unweighted UniFrac distance. C. Each bar represents the mean ± SEM. D. PCoA of Bray-Curtis distance. E. Each bar represents the mean ± SEM. \**P <* 0.05, \*\**P <* 0.01. According to non-parametric Kruskal-Wallis test with Bonferroni post hoc and one-way ANOVA. n = 7–8.

Principal coordinates analysis (PCoA) of the unweighted UniFrac distance matrix showed that the gut microbial composition differed significantly among the three groups and that soil intervention contributed to the unique microbial structure and composition (Fig. 1B). The distance between the Soil group and Asthma group or PBS group was significantly greater than that between the PBS and Asthma groups (Fig. 1C and Table E2), suggesting the similarity between the Asthma and PBS groups was the highest, while the difference between the intestinal microorganisms in the Soil group and Asthma and PBS groups was more pronounced, indicating that soil significantly affected the gut microbial composition. Furthermore, the PCoA of the Bray–Curtis matrix was similar to that of the unweighted UniFrac distance matrix (Fig. 1D and 1E).

At the phylum level, the gut microbial communities of all three groups mice were dominated by Bacteroidetes and Firmicutes (Fig. E3B and Table E3), and the Bacteroidetes/Firmicutes ratio of the Soil and Asthma groups was significantly lower than that of the PBS group, but no significant differences were found between the Asthma and Soil groups (Fig. E3C). Compared to the Asthma group, the abundance of Actinobacteria was significantly lower in both PBS and Soil groups (Fig. 2A). Significant changes were found in the genus composition of Actinobacteria (Fig. 2B), Firmicutes (Fig. 2C), and Bacteroidetes (Fig. E3D). Soil group mice showed a significant change in the percentage of genus composition with a higher abundance of *Allobaculum*, *Alistipes*, *Lachnospiraceae_UCG-001*, *Rikenellaceae_RC9_gut_group*, and *Christensenellaceae_R-7_group*, and soil intake decreased *Turicibacter* and *Eubacterium-xylanophilum-group* abundance (Fig. 2D, Fig. E3E and Table E4). The heatmap of the top 250 OTUs revealed that soil intake had a significant effect on OTUs in asthmatic mice and that it promoted the growth of some bacterial OTUs (Fig. 2E).

**Figure 2.**
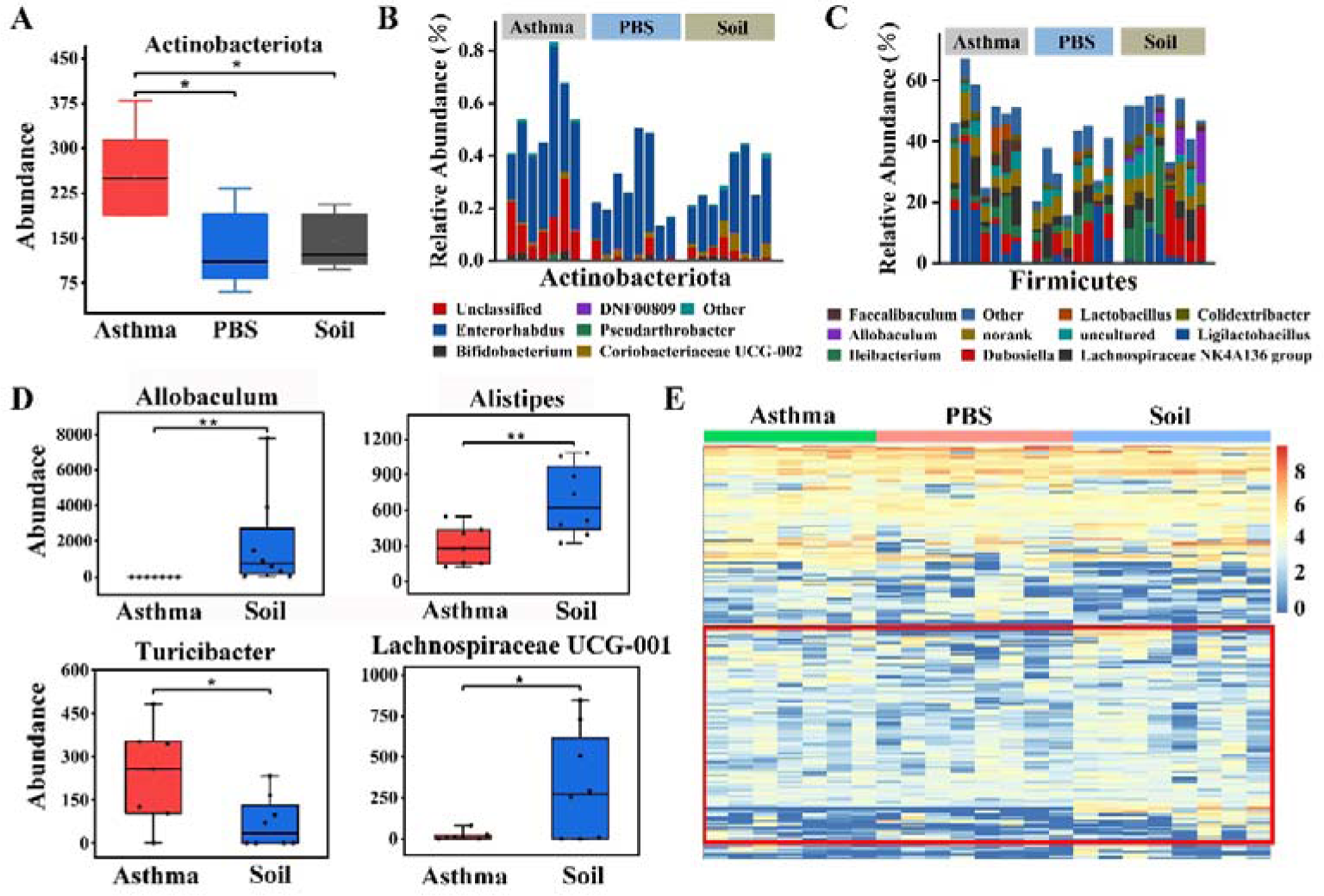
Comparison of intestinal microbial composition and structure. A. Actinobacteriota abundance. B, C. Detailed relative abundance of bacterial genera in phyla Actinobacteriota and Firmicutes. D. Abundance of bacterial genera between Asthma and Soil groups. E. Heatmap for the impact of sterilized soil intake on OTUs at a 97% threshold. \**P <* 0.05, \*\**P <* 0.01. According to two-tailed least significant difference test. n = 7–8.

Linear discriminant analysis effect size (LEfSe) analysis was performed to identify the most differentially abundant taxa between Asthma and Soil groups. LEfSe analysis showed a significant increase in the phylum Actinobacteria in Asthma group compared to that in Soil group. Furthermore, *Allobaculum*, *Alistipes* (Fig. 3A and 3B), and *Rikenellaceae-RC9-gut-group* were significantly enriched in Soil group while *Eubacterium-xylanophilum-group* and *Turicibacter* (Fig. E3F) dominated in Asthma group mice. To reveal the relationship between samples and bacterial abundance, circos analysis was performed. Circos plot demonstrated that *Lachnospiraceae_NK4A136_group*, *lleibacterium,* and *Allobaculum* were more abundant in the Soil group, while the association with *Ligilactobacillus* appeared to be clearer in the Asthma group (Fig. 3C). Interestingly, the growth-promoting bacteria with soil, including *Allobaculum*, *Alistipes*, *Lachnospiraceae_UCG-001, Rikenellaceae_RC9_gut_group,* and *Christensenellaceae_R-7_group,* could produce short-chain fatty acids (SCFA)(19–23), belonging to beneficial bacteria in the intestine while *Turicibacter*, whose growth is inhibited, was previously shown to be a conditionally pathogenic bacterium(24) and the core genus in HDM-induced asthma model(25).

**Figure 3.**
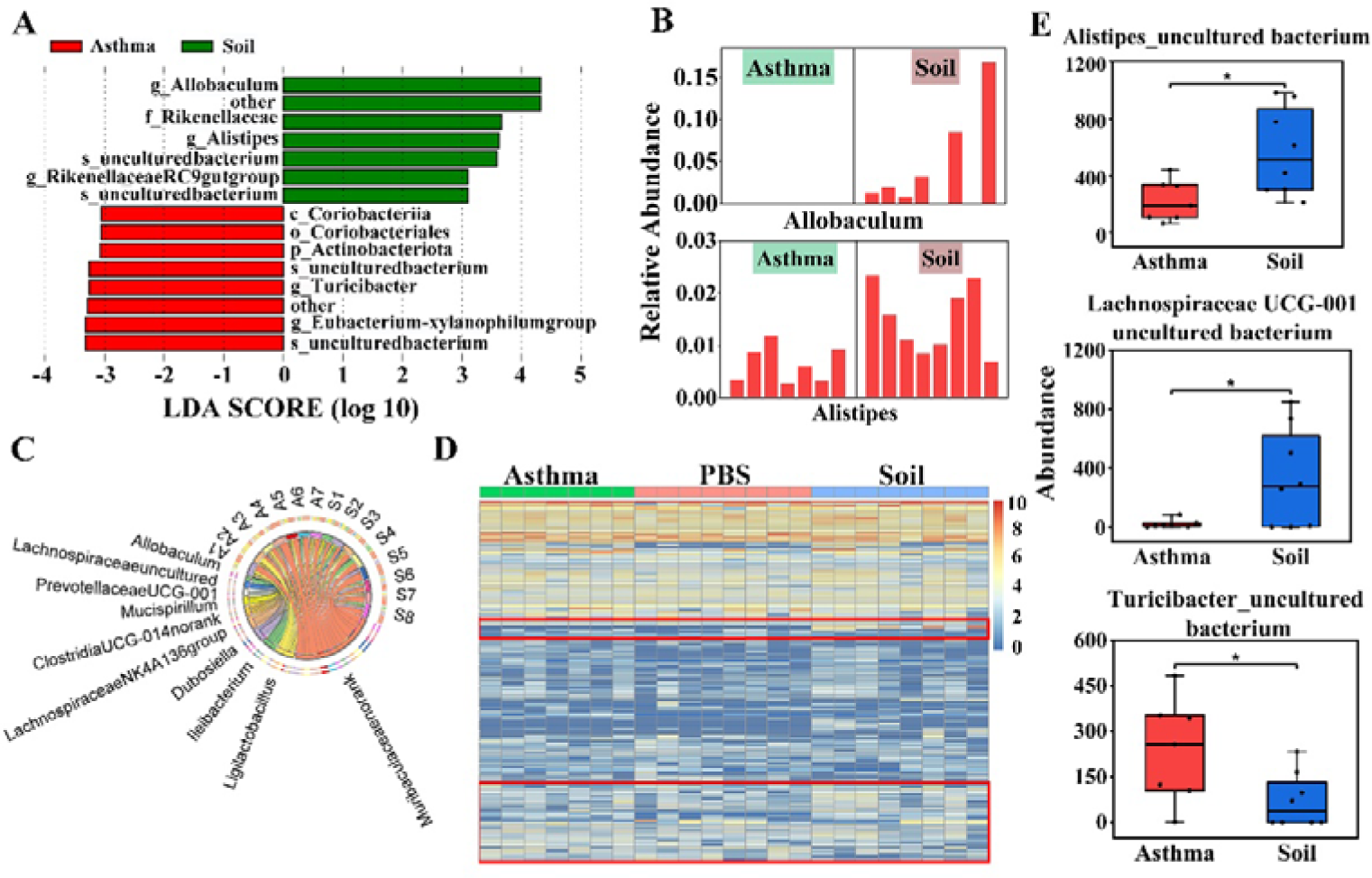
Soil intake effects on OTUs, genera, and species in asthmatic mice. A. LEfSe identified the most differentially abundant taxa between Asthma and Soil groups at the genus level. Soil-enriched taxa are indicated with a positive LDA score (green) and taxa enriched in the Asthma group with a negative score (red). Only taxa meeting an LDA significant threshold of > 3.0 are shown. B. Relative abundance of biomarkers identified by LFfSe in mice of each group. C. Circos plot. The left semi-circle was the distribution of the more dominant bacterial genus in each sample, and each bacterial genus was assigned a specific color. The right semi-circle represents different samples. S: Soil; A: Asthma. D. Heatmap for the impact of sterilized soil intake on species level. E. Differential species between the two groups.\**P <* 0.05, \*\**P <* 0.01. According to two-tailed least significant difference test. n = 7–8.

Heatmap analysis on the species level revealed that the species composition of the intestinal bacterial community changed after soil intake (Fig. 3D). Specifically, soil ingestion boosted the growth of *Lachnospiraceae UCG-001_uncultured bacterium, Alistipes_uncultured bacterium, Rikenellaceae RC9 gut group_uncultured bacterium, Parabacteroides_Unclassified,* and *Alistipes_Unclassified*, whereas the abundance of *[Eubacterium] xylanophilum group_uncultured bacterium, Turicibacter_uncultured bacterium, Ligilactobacillus_Unclassified,* and *Atopobiaceae_Unclassified* decreased in the Soil group (Fig. 3E, Fig. E3G and Table E5).

Hence, soil intake significantly altered the mouse gut microbial structure and composition, and promoted the growth of various beneficial bacteria. OVA sensitization also significantly affected the mouse intestinal microbial composition, but the effect was not as strong as that of soil intervention.

### Soil intake attenuated lung inflammation and alleviated Th2-type allergic responses

To study the therapeutic effect of soil intake on OVA-induced lung inflammation, we examined the immune status of the lung, BALF, and serum. Hematoxylin and eosin (HE) and periodic acid-Schiff (PAS) staining results showed that the alveolar structure was intact in the PBS group, with no evident inflammatory cell infiltration or mucus in the lung. In contrast, Asthma group mice developed severe inflammation with more inflammatory cell infiltration and lung tissue mucus, while soil ingestion significantly reduced the numbers of total cells, eosinophils, and neutrophils, and lung inflammation was improved in Soil group (Fig. 4A and 4B).

**Figure. 4.**
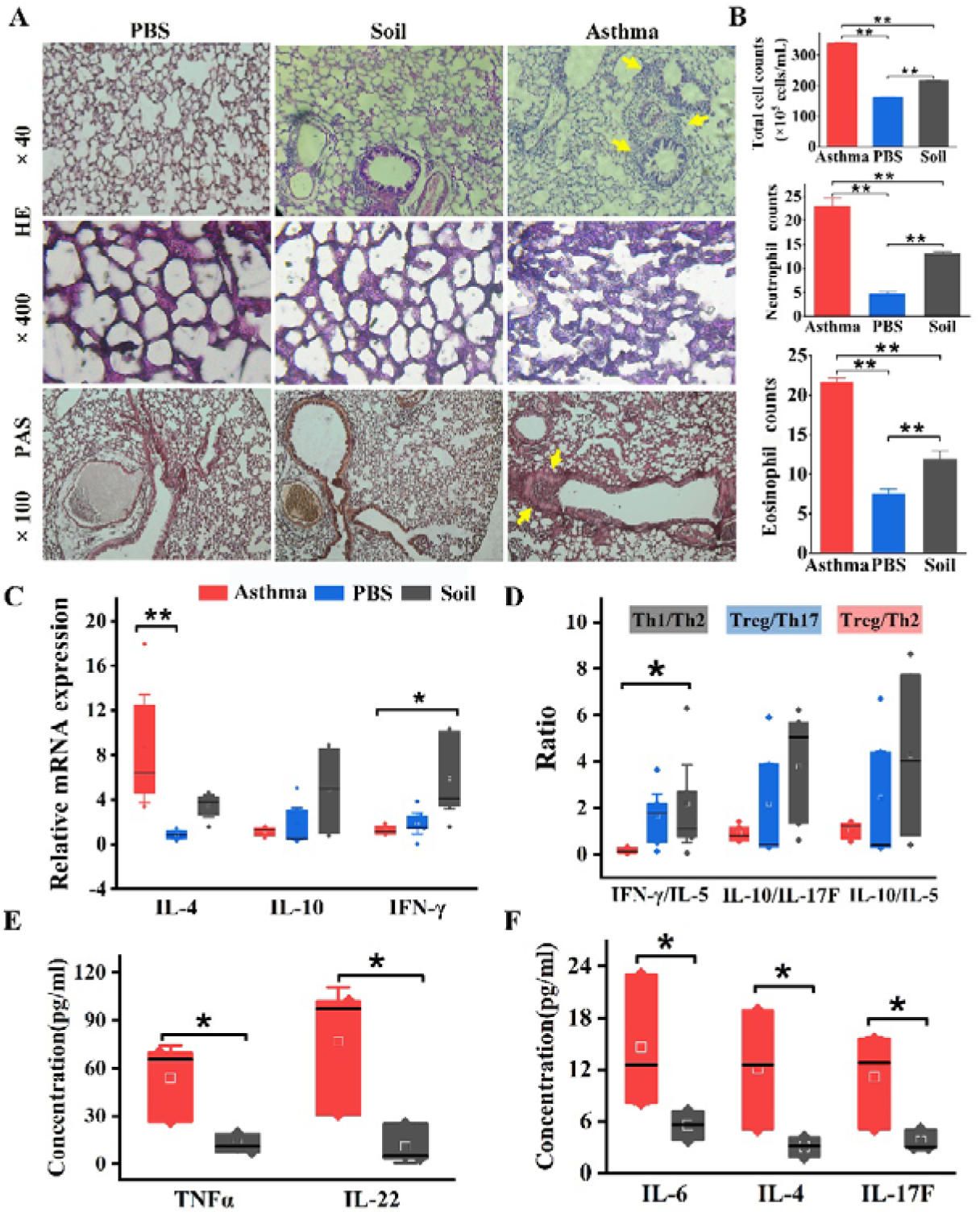
Soil intake attenuated allergic asthma symptoms. A. Representative micrographs of HE– and PAS-stained lung tissue. Glycogen and other PAS-positive substances in the tissue were conspicuously red in PAS. B. Numbers of the total cell, eosinophils, and neutrophils in BALF. A total of 200 cells were counted per slide when counting eosinophils and neutrophils. C, D. Concentrations of TNF-α, IL-22, IL-6, IL-4 and IL-17F in serum. E. Lung mRNA levels of IL-4, IL-10, and IFN-γ. F. Effects of soil ingestion on the balance of Th1/Th2, Treg/Th17 and Treg/Th2. *\*P* < 0.05, *\*\*P* < 0.01. According to one-way ANOVA and the non-parametric Mann-Whitney U test and Kruskal-Wallis test. n = 7–8.

To further explore the underlying mechanisms, the gene expression levels of asthma-related cytokines in the lung were investigated. Soil intake upregulated IFN-γ and downregulated IL-4 in the lung, which altered the Th1/Th2 balance and skewed the immune system toward a higher level of anti-inflammatory signaling, attenuating OVA-induced allergic asthma responses (Fig. 4C, 4D and Table E6). IL-10 levels in the lung were relatively higher than those of IL-17F and IL-5 in the Soil group compared with the PBS and Asthma groups, though this difference was not significant (Fig. 4D and Table E6). Furthermore, we examined the changes in serum immune parameters. Soil intake decreased the serum concentrations of IL-4, IL-6, IL-17F, TNF-α, and IL-22, cytokines, which can elevate the asthma exacerbation risk (Fig. 4E and 4F).

Thus, soil intervention had a significant therapeutic effect on mouse asthma. Reduced Th2-type allergic response was one of the underlying mechanisms. In contrast, no significant effect on asthma was found with simple air microorganisms exposure in the general animal rooms.

### Influence of OVA sensitization, air microbes, and age on the gut microbial composition

The “gut-lung axis” states that sensitization processes contribute to changes in intestinal microorganisms, and that gut microbial dysbiosis can in turn aggravate lung inflammation(26). To investigate the effect of OVA sensitization on intestinal microbial composition in mice, we analyzed fecal microbiota. PCoA based on the unweighted UniFrac distance showed obvious differences between the P_Asthma and P_PBS groups (Fig. 5A). Furthermore, nonmetric multidimensional scaling analysis was similar to that of the unweighted UniFrac distance matrix (Fig. 5B).

**Figure. 5.**
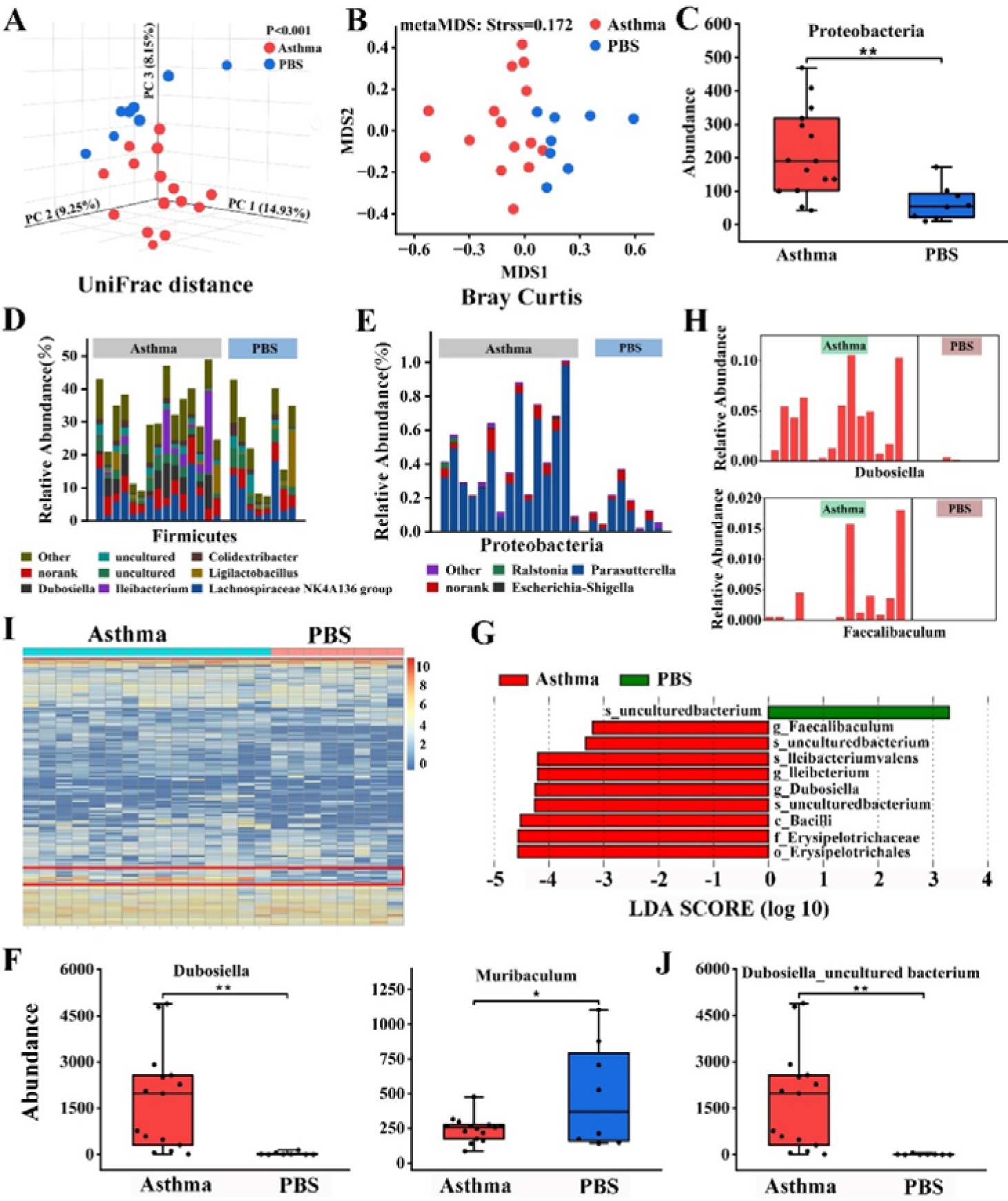
Sensitization altered gut microbial composition. A. PCoA of unweighted UniFrac distance. B. Nonmetric multidimensional scaling of unweighted UniFrac distance. C. The abundance of fecal bacterial phylum Proteobacteria. D, E. Detailed relative abundance of bacterial genera in phyla Firmicutes and Proteobacteria. F. Differential genera. G. LEfSe analysis for characteristic microbial genera in P_Asthma and P_PBS groups with an LDA score > 3.2. H. Relative abundance of the most differentially abundant genera identified by LFfSe. I. Heatmap at species level. J. Differential species. \**P <* 0.05, \*\**P <* 0.01. According to two-tailed least significant difference test. n = 8–2.

OVA sensitization changed the microbial communities at the phylum level, with a lower abundance of Proteobacteria (Fig. 5C and Table E7). Both the gut microbial communities of P_Asthma and P_PBS mice were dominated by the Bacteroidetes and Firmicutes (Fig. E4a), and sensitization did not change Bacteroidetes/Firmicutes ratio (Fig. E4B). Detailed genera compositions of Firmicutes (Fig. 5D), Proteobacteria (Fig. 5E), and Bacteroidata (Fig. E4C) were shown. At the genus level, a higher abundance of *Dubosiell*, *Parasutterell*, *Rikenella*, and *Catenibacterium* was observed in P_Asthma group. In contrast, the abundance of *Muribaculum, Staphylococcus,* and *Butyricicoccus* decreased in asthmatic mice (Fig. 5F, Fig. E4D and Table E8). LEfSe results identified three bacterial genera (*Dubosiella*, *Ileibacterium*, and *Faecalibaculum*) enriched in the P_Asthma group with an LDA score > 3.2 (Fig. 5G, 5H and Fig. E4F). Heatmap on the species level revealed that sensitization changed species composition of the intestinal bacterial community (Fig. 5I), and sensitization promoted the growth of *Dubosiella_uncultured bacterium*, *Burkholderiales bacterium YL45*, *Bacteroides_uncultured bacterium*, and *Catenibacterium_uncultured bacterium*, while the abundance of *Muribaculum_uncultured bacterium*, *Staphylococcus_Unclassified*, and *Butyricicoccus_uncultured bacterium* decreased in asthmatic mice (Fig. 5J, Fig. E4E and Table E9). Thus, sensitization has a significant effect on gut microbial composition and structure.

Several reports indicate that farm environmental microbes(4,27) and age(28,29) affect the gut microbiota. We investigated these findings, and the PCoA results (Fig. E5A) showed a significant difference between the P-Asthma and Asthma groups, and between the P-PBS and PBS groups. However, the effect of air microorganisms and age on mouse gut microbiota is weaker than that of OVA sensitization (Fig. E5B), and largely weaker than that of soil (Fig. E6). The LEfSe results showed (Fig. E7) that for asthmatic mice, air microbes and age significantly increased the phylum Firmicutes, and genera *Ligilactobacillus* and *Faecalibaculum*, while for normal mice, air microbes and age significantly increased the genera *Dubosiella*, *Ileibacterium,* and *Lactobacillus* (Fig. E8).

In conclusion, air microorganisms and age, and sensitization had significant effects on mouse gut microbes, with sensitization having greater effects on mouse gut microbes than air microorganisms and age together, but all these effects were weaker than those of soil.

## Discussion

We used OVA-induced asthma model mice to investigate the effects of sterilized soil intake, as a prebiotic, accompanied by air microorganisms exposure, on asthmatic mice. We found that soil prebiotics significantly altered the gut microbial composition and exerted significant therapeutic effects on asthmatic mice. Further studies revealed that this therapeutic effect was achieved by reducing the serum concentrations of Th2-related cytokines and regulating the Th1/Th2 balance by enhancing anti-inflammatory signaling in the lung.

Soil has a therapeutic effect on asthma, possibly by increasing the growth of a variety of beneficial commensal bacteria that are protective against allergies in the intestine, including *Alistipes*, *Rikenellaceae_RC9_gut group*, *Allobaculum*, *Ileibacterium,* and *Christensenellaceae* and decreasing pathogenic bacteria abundance associated with asthma. Several studies have shown that a reduced abundance of the *Alistipes* was observed in populations with asthma and food allergy(30,31) and a higher abundance of the *Alistipes* correlated with higher fecal and serum acetate and fecal butyrate levels(32). Similarly, the *Rikenellaceae_RC9_gut group* is positively correlated with butyric and valeric acid and can alleviate colitis symptoms by stimulating Treg cell differentiation(33,34). *Lachnospiraceae UCG-001*(35) and *Ileibacterium*(36), which can produce SCFA, have been reported to be beneficial against inflammation. *Allobaculum* was related to the increase in SCFA production, especially butyrate(19,20,37). *Christensenellaceae* is widespread in humans and animals, playing a key role in maintaining host health(38).

The effects of soil intake on mouse asthma, regardless of therapeutic effects or mechanism, are consistent with those of LCLE exposure. Firstly, it can be seen from the HE staining and BALF that soil intake and keeping mice in an LCLE(6) have similar protective effects against lung inflammation. Further, asthma was prevented in mice housed in cages with soil with lower expression of IL-5 and IL-13 in lung and higher expression of IFN-γ, altering the Th1/Th2 balance(6). Similarly, our results showed that the IFN-γ level was significantly higher in lung, which altered the Th1/Th2 balance, skewing the immune system toward higher levels of anti-inflammatory signaling, and thereby attenuating allergic responses. Exposure to forests, grasslands, climbing, and digging mud can increase the serum IL-10:IL-17A ratio among daycare children(39). Soil intake also affected the Treg/Th17 balance, and the expression of anti-inflammatory IL-10 relative to that of IL-5 in the lung was greater in Soil group, which positively improved asthma symptoms although the difference was not significant. González-Rodríguez et al. found that exposure to dust derived from sterilized forest soil reduced the levels of IL-17F in serum and TNF-α in splenocytes(40). Similarly, in our study, significant reductions in the serum concentrations of IL-17F and TNF-α were observed after soil ingestion. Furthermore, the concentrations of serum IL-6 and IL-22 were significantly lower after soil intake. Increased serum IL-6 level was observed in asthmatic patients and correlated with the risk of asthma exacerbation(41,42). IL-22, an important effector molecule produced by Th17 cells, can drive airway hyperresponsiveness in combination with IL-17(43). The reduction in the numbers of total cells, eosinophils, and neutrophils in BALF in Soil group further supported the alleviation of asthma symptoms. Inflammatory cell infiltration in lung was significantly reduced, the alveolar wall structure was more intact, and mucus secretion in the bronchi was considerably improved after soil intervention.

Although edible soil and LCLE exposure can effectively prevent or treat allergic asthma, edible soil has more advantages than LCLE treatment: (I) In contrast to LCLE, sterilized soil does not contain pathogens causing infections; (II) LCLE does not conform to the lifestyle of modern people; (III) taking soil as a prebiotic to treat asthma is more convenient; and (IV) edible soil is safe. Since pre-history, humans have taken mineral– and trace element-rich soil, and used selected soils for medicinal purposes, usually for the treatment of gastrointestinal ailments(44). Some commercial medicines (e.g., Kaopectate), contain soil components and are widely used to treat diarrhea, nausea, and other diseases. Moreover, soil contains a large number of natural clay minerals, such as montmorillonite and kaolinite(45), which are traditional Chinese medicines used to treat certain diseases. In addition, heat-killed bacteria(46,47) and bacterial endotoxin(48) can also effectively stimulate our immune response. Further study investigating the reasons for the immune changes caused by soil ingestion is warranted.

Soil ingestion altered the gut microbial composition in asthmatic mice, but no increase in intestinal microbial diversity or richness was found, which is inconsistent with the findings by Zhou et al.(16). Presumably, this may be due to the influence of stress. In this experiment, mice were challenged with OVA in a confined space for 40 min to induce asthma symptoms and our sampling time was 24 h post-challenge. The effects of stress on the gut microbiota have been previously reported(49,50) and gut microbial diversity needs sufficient time to recover(8). Ottman et al. did not observe changes in intestinal microbial diversity when the asthma model was established in mice raised in LCLE(6), consistent with our results.

It is unclear how soil supports the growth of a variety of beneficial commensal bacteria. Presumably, natural clay minerals that are widespread in soil, including montmorillonite, kaolinite, and illite, may play a key role in shaping the gut microbiota. These clay minerals have a large specific surface area and cation exchange capacity per unit mass, so they can adsorb microbial cells and obtain nutrients (e.g., polysaccharides secreted by microorganisms) to interact with soil microorganisms to affect their growth and function and support biofilm formation(45), which is key to improving bacterial species colonization. Oral administration of montmorillonite can help probiotics, especially lactic acid bacteria (LABs), to form biofilm on its surface and further promote the growth of LABs to enhance anti-tumor responses(51). The gut is a large reservoir inhabiting trillions of bacteria. Hence, we speculate these microbes can interact extensively with clay minerals in soil, and microorganisms in the intestine may form biofilms on the surface of these particles to modify the gut microbiota, which might be a potential mechanism of soil-induced changes in intestinal microbes.

In our study, both sensitization and air microorganisms and age had a significant effect on mouse intestinal microbes. Two possible factors affect the intestinal microorganisms during sensitization, the sensitization-caused inflammatory state and stress responses. Both the inflammatory state(52) and stress(49) affect the mouse gut microbes, consistent with our results. However, further experiments are needed to prove which factor is more influential. Furthermore, air microorganisms and age also significantly affected the intestinal microorganisms in mice. Both air microorganisms(4) and age(28) had an effect on gut microorganisms, but in this study, the effect of age may be smaller because the samples were collected twice in the adult stage, and the microbiota display a stable homeostatic state in adulthood(53). Thus, air microorganisms may have a greater effect on mouse gut microorganisms than age, but their effects are significantly smaller than those of soil. Surprisingly, the effect of sensitization on gut microbes was greater than that of air microbes and age together.

## Conclusion

In conclusion, soil intake had a significant therapeutic effect on mouse asthma, achieved by promoting the growth of multiple beneficial bacteria. The protective effects and mechanism of soil intake on asthma are consistent with those of LCLE, but edible soil is easier to implement in the clinic. The results indicated that the development of soil-based prebiotic products might be used for allergic asthma intervention.

## Acknowledgments

This work was supported by The Natural Science Foundation of China (grant no. 31770540), The Key Research Program of Jiangsu (grant no. BE2018663) and the Fundamental Research Funds for the Central Universities (grant nos. 2242021k30014 and 2242021k30059).

We would like to thank Editage (www.editage.cn) for English language editing.

## References

1. Lin CH. Treatment of Hypertension in Patients with Asthma. N Engl J Med. 2019 Dec 5;381(23):2278–9.

2. Ege MJ, Mayer M, Normand AC, Genuneit J, Cookson WOCM, Braun-Fahrländer C, et al. Exposure to environmental microorganisms and childhood asthma. N Engl J Med. 2011 Feb 24;364(8):701–9.

3. Stein MM, Hrusch CL, Gozdz J, Igartua C, Pivniouk V, Murray SE, et al. Innate Immunity and Asthma Risk in Amish and Hutterite Farm Children. N Engl J Med. 2016 Aug 4;375(5):411–21.

4. Depner M, Taft DH, Kirjavainen PV, Kalanetra KM, Karvonen AM, Peschel S, et al. Maturation of the gut microbiome during the first year of life contributes to the protective farm effect on childhood asthma. Nat Med. 2020 Nov;26(11):1766–75.

5. Zhou D, Zhang H, Bai Z, Zhang A, Bai F, Luo X, et al. Exposure to soil, house dust and decaying plants increases gut microbial diversity and decreases serum immunoglobulin E levels in BALB/c mice. Environ Microbiol. 2016 May;18(5):1326–37.

6. Ottman N, Ruokolainen L, Suomalainen A, Sinkko H, Karisola P, Lehtimäki J, et al. Soil exposure modifies the gut microbiota and supports immune tolerance in a mouse model. J Allergy Clin Immunol. 2019 Mar;143(3):1198–1206.e12.

7. Vo N, Tsai TC, Maxwell C, Carbonero F. Early exposure to agricultural soil accelerates the maturation of the early-life pig gut microbiota. Anaerobe. 2017 Jun;45:31–9.

8. Li N, Zhang H, Bai Z, Jiang H, Yang F, Sun X, et al. Soil exposure accelerates recovery of the gut microbiota in antibiotic-treated mice. Environ Microbiol Rep. 2021 Oct;13(5):616–25.

9. Nurminen N, Lin J, Grönroos M, Puhakka R, Kramna L, Vari HK, et al. Nature-derived microbiota exposure as a novel immunomodulatory approach. Future Microbiol. 2018 Jun 1;13:737–44.

10. Forsythe P, Inman MD, Bienenstock J. Oral treatment with live Lactobacillus reuteri inhibits the allergic airway response in mice. Am J Respir Crit Care Med. 2007 Mar 15;175(6):561–9.

11. Huang CF, Chie WC, Wang IJ. Efficacy of Lactobacillus Administration in School-Age Children with Asthma: A Randomized, Placebo-Controlled Trial. Nutrients. 2018 Nov 5;10(11):E1678.

12. Wawryk-Gawda E, Markut-Miotła E, Emeryk A. Postnatal probiotics administration does not prevent asthma in children, but using prebiotics or synbiotics may be the effective potential strategies to decrease the frequency of asthma in high-risk children – a meta-analysis of clinical trials. Allergol Immunopathol (Madr). 2021;49(4):4–14.

13. Wu Y, Chen Y, Li Q, Ye X, Guo X, Sun L, et al. Tetrahydrocurcumin alleviates allergic airway inflammation in asthmatic mice by modulating the gut microbiota. Food Funct. 2021 Aug 2;12(15):6830–40.

14. Kang Y, Cai Y. Future prospect of faecal microbiota transplantation as a potential therapy in asthma. Allergol Immunopathol (Madr). 2018 Jun;46(3):307–9.

15. Zhou D, Bai Z, Zhang H, Li N, Lu Z. Soil is a key factor influencing gut microbiota and its effect is comparable to that exerted by diet for mice. F1000 Res. 2018;7:1588.

16. Zhou D, Li N, Yang F, Zhang H, Bai Z, Dong Y, et al. Soil causes gut microbiota to flourish and total serum IgE levels to decrease in mice. Environ Microbiol. 2022 Mar 22;

17. Lathrop SK, Bloom SM, Rao SM, Nutsch K, Lio CW, Santacruz N, et al. Peripheral education of the immune system by colonic commensal microbiota. Nature. 2011 Sep 21;478(7368):250–4.

18. Grigg JB, Sonnenberg GF. Host-Microbiota Interactions Shape Local and Systemic Inflammatory Diseases. J Immunol Baltim Md 1950. 2017 Jan 15;198(2):564–71.

19. Zhang X, Zhao Y, Zhang M, Pang X, Xu J, Kang C, et al. Structural changes of gut microbiota during berberine-mediated prevention of obesity and insulin resistance in high-fat diet-fed rats. PloS One. 2012;7(8):e42529.

20. Wu M, Yang S, Wang S, Cao Y, Zhao R, Li X, et al. Effect of Berberine on Atherosclerosis and Gut Microbiota Modulation and Their Correlation in High-Fat Diet-Fed ApoE-/– Mice. Front Pharmacol. 2020;11:223.

21. Guo M, Li Z. Polysaccharides isolated from Nostoc commune Vaucher inhibit colitis-associated colon tumorigenesis in mice and modulate gut microbiota. Food Funct. 2019 Oct 16;10(10):6873–81.

22. Holman DB, Gzyl KE. A meta-analysis of the bovine gastrointestinal tract microbiota. FEMS Microbiol Ecol. 2019 Jun 1;95(6):fiz072.

23. López-Montoya P, Cerqueda-García D, Rodríguez-Flores M, López-Contreras B, Villamil-Ramírez H, Morán-Ramos S, et al. Association of Gut Microbiota with Atherogenic Dyslipidemia, and Its Impact on Serum Lipid Levels after Bariatric Surgery. Nutrients. 2022 Aug 28;14(17):3545.

24. Gangadoo S, Dinev I, Chapman J, Hughes RJ, Van TTH, Moore RJ, et al. Selenium nanoparticles in poultry feed modify gut microbiota and increase abundance of Faecalibacterium prausnitzii. Appl Microbiol Biotechnol. 2018 Feb;102(3):1455–66.

25. Hu W, Lu W, Li L, Zhang H, Lee YK, Chen W, et al. Both living and dead Faecalibacterium prausnitzii alleviate house dust mite-induced allergic asthma through the modulation of gut microbiota and short-chain fatty acid production. J Sci Food Agric. 2021 Oct;101(13):5563–73.

26. Penders J, Stobberingh EE, van den Brandt PA, Thijs C. The role of the intestinal microbiota in the development of atopic disorders. Allergy. 2007 Nov;62(11):1223–36.

27. Shimojo N, Izuhara K. Old friends, microbes, and allergic diseases. Allergol Int Off J Jpn Soc Allergol. 2017 Oct;66(4):513–4.

28. Yatsunenko T, Rey FE, Manary MJ, Trehan I, Dominguez-Bello MG, Contreras M, et al. Human gut microbiome viewed across age and geography. Nature. 2012 May 9;486(7402):222–7.

29. O’Toole PW, Jeffery IB. Gut microbiota and aging. Science. 2015 Dec 4;350(6265):1214–5.

30. Zolnikova OY, Potskhverashvili ND, Kudryavtseva AV, Krasnov GS, Guvatova ZG, Truhmanov AS, et al. [Changes in gut microbiota with bronchial asthma]. Ter Arkh. 2020 Apr 27;92(3):56–60.

31. Chen CC, Chen KJ, Kong MS, Chang HJ, Huang JL. Alterations in the gut microbiotas of children with food sensitization in early life. Pediatr Allergy Immunol Off Publ Eur Soc Pediatr Allergy Immunol. 2016 May;27(3):254–62.

32. Medawar E, Haange SB, Rolle-Kampczyk U, Engelmann B, Dietrich A, Thieleking R, et al. Gut microbiota link dietary fiber intake and short-chain fatty acid metabolism with eating behavior. Transl Psychiatry. 2021 Oct 1;11(1):500.

33. Qing Y, Xie H, Su C, Wang Y, Yu Q, Pang Q, et al. Gut Microbiome, Short-Chain Fatty Acids, and Mucosa Injury in Young Adults with Human Immunodeficiency Virus Infection. Dig Dis Sci. 2019 Jul;64(7):1830–43.

34. Dubin K, Callahan MK, Ren B, Khanin R, Viale A, Ling L, et al. Intestinal microbiome analyses identify melanoma patients at risk for checkpoint-blockade-induced colitis. Nat Commun. 2016 Feb 2;7:10391.

35. Wei X, Tao J, Xiao S, Jiang S, Shang E, Zhu Z, et al. Xiexin Tang improves the symptom of type 2 diabetic rats by modulation of the gut microbiota. Sci Rep. 2018 Feb 27;8:3685.

36. Wang Y, Ablimit N, Zhang Y, Li J, Wang X, Liu J, et al. Novel β-mannanase/GLP-1 fusion peptide high effectively ameliorates obesity in a mouse model by modifying balance of gut microbiota. Int J Biol Macromol. 2021 Nov 30;191:753–63.

37. Balakrishnan B, Luckey D, Bodhke R, Chen J, Marietta E, Jeraldo P, et al. Prevotella histicola Protects From Arthritis by Expansion of Allobaculum and Augmenting Butyrate Production in Humanized Mice. Front Immunol. 2021;12:609644.

38. Waters JL, Ley RE. The human gut bacteria Christensenellaceae are widespread, heritable, and associated with health. BMC Biol. 2019 Oct 28;17(1):83.

39. Roslund MI, Puhakka R, Grönroos M, Nurminen N, Oikarinen S, Gazali AM, et al. Biodiversity intervention enhances immune regulation and health-associated commensal microbiota among daycare children. Sci Adv. 2020 Oct;6(42):eaba2578.

40. González-Rodríguez MI, Nurminen N, Kummola L, Laitinen OH, Oikarinen S, Parajuli A, et al. Effect of inactivated nature-derived microbial composition on mouse immune system. Immun Inflamm Dis. 2022 Mar;10(3):e579.

41. Yokoyama A, Kohno N, Fujino S, Hamada H, Inoue Y, Fujioka S, et al. Circulating interleukin-6 levels in patients with bronchial asthma. Am J Respir Crit Care Med. 1995 May;151(5):1354–8.

42. Jackson DJ, Bacharier LB, Calatroni A, Gill MA, Hu J, Liu AH, et al. Serum IL-6: A biomarker in childhood asthma? J Allergy Clin Immunol. 2020 Jun;145(6):1701–1704.e3.

43. Whitehead GS, Kang HS, Thomas SY, Medvedev A, Karcz TP, Izumi G, et al. Therapeutic suppression of pulmonary neutrophilia and allergic airway hyperresponsiveness by a RORγt inverse agonist. JCI Insight. 2019 Jun 11;5:125528.

44. Sing D, Sing CF. Impact of direct soil exposures from airborne dust and geophagy on human health. Int J Environ Res Public Health. 2010 Mar;7(3):1205–23.

45. Zhang L, Gadd GM, Li Z. Microbial biomodification of clay minerals. Adv Appl Microbiol. 2021;114:111–39.

46. Park S, Kim JI, Bae JY, Yoo K, Kim H, Kim IH, et al. Effects of heat-killed Lactobacillus plantarum against influenza viruses in mice. J Microbiol Seoul Korea. 2018 Feb;56(2):145–9.

47. Miyazawa K, Kawase M, Kubota A, Yoda K, Harata G, Hosoda M, et al. Heat-killed Lactobacillus gasseri can enhance immunity in the elderly in a double-blind, placebo-controlled clinical study. Benef Microbes. 2015;6(4):441–9.

48. Schuijs MJ, Willart MA, Vergote K, Gras D, Deswarte K, Ege MJ, et al. Farm dust and endotoxin protect against allergy through A20 induction in lung epithelial cells. Science. 2015 Sep 4;349(6252):1106–10.

49. K V, K SP, M H, B P, Kj P, A T, et al. Impact of stress on the gut microbiome of free-ranging western lowland gorillas. Microbiol Read Engl [Internet]. 2018 Jan [cited 2022 Aug 1];164(1). Available from: https://pubmed.ncbi.nlm.nih.gov/29205130/

50. Bailey MT, Dowd SE, Parry NMA, Galley JD, Schauer DB, Lyte M. Stressor exposure disrupts commensal microbial populations in the intestines and leads to increased colonization by Citrobacter rodentium. Infect Immun. 2010 Apr;78(4):1509–19.

51. Han C, Song J, Hu J, Fu H, Feng Y, Mu R, et al. Smectite promotes probiotic biofilm formation in the gut for cancer immunotherapy. Cell Rep. 2021 Feb 9;34(6):108706.

52. Arrieta MC, Stiemsma LT, Dimitriu PA, Thorson L, Russell S, Yurist-Doutsch S, et al. Early infancy microbial and metabolic alterations affect risk of childhood asthma. Sci Transl Med. 2015 Sep 30;7(307):307ra152.

53. Laukens D, Brinkman BM, Raes J, De Vos M, Vandenabeele P. Heterogeneity of the gut microbiome in mice: guidelines for optimizing experimental design. FEMS Microbiol Rev. 2016 Jan;40(1):117–32.

