## Supplementary Figures and Tables for "Soil intake modifies the gut microbiota and alleviates ovalbumin-induced mice asthma inflammation"

**Figure E1**

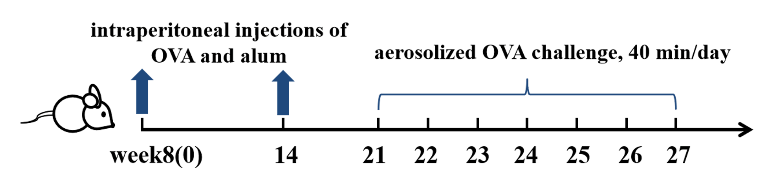

**Figure E1. OVA-induced asthma model protocol.** Eight-week-old mice were injected with the mixture of OVA and alum on day 0 and day 14 respectively and received an OVA challenge of 40 min/day from day 21 to day 27.

**Figure E2**

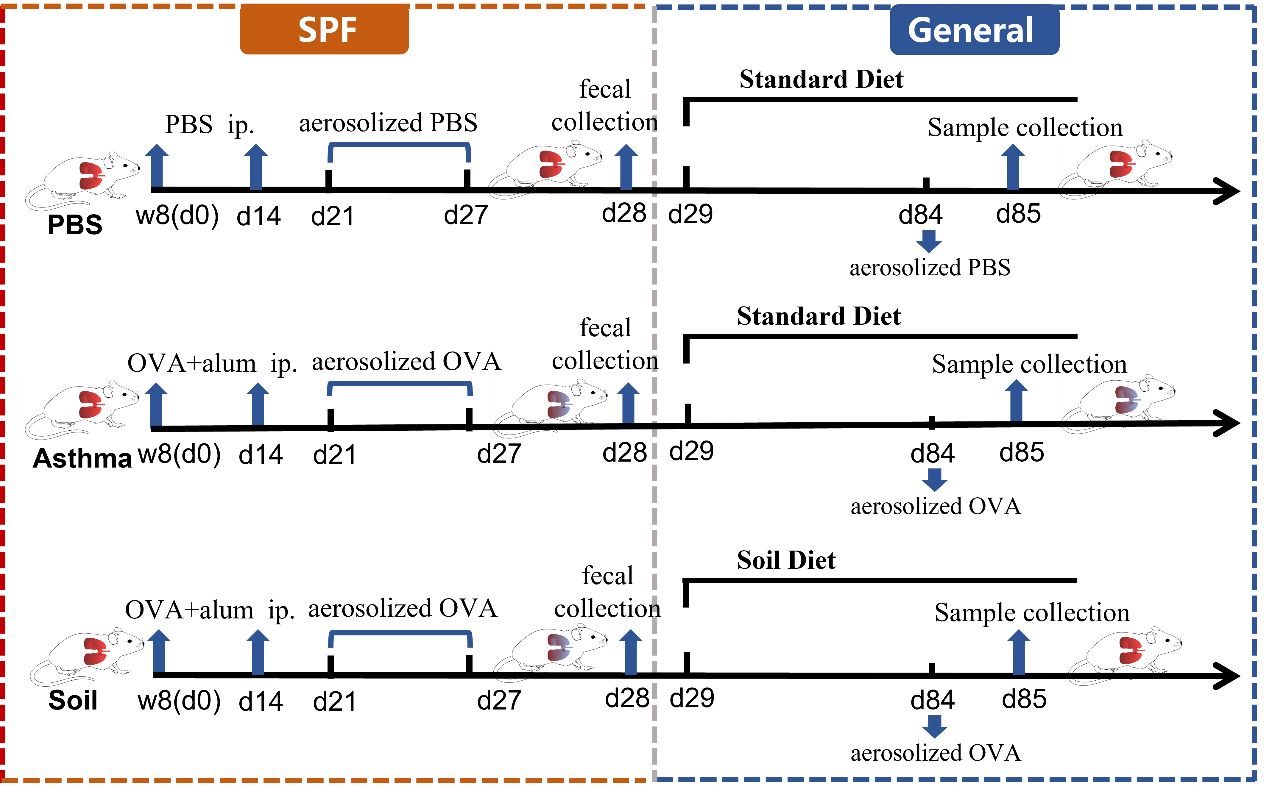

**Figure E2. Study design timeline.** In the SPF animal facility, eight-week-old SPF mice were intraperitoneally injected with OVA and alum adjuvant on days 0 and 14, respectively, followed by aerosolized OVA challenge to induce an asthma model. Control group mice were treated with PBS and mouse feces were collected on day 28. All mice were, then, transferred to the general animal room. Asthma and PBS group mice were provided with a standard diet, while Soil group mice consumed a standard diet containing 5% sterilized soil collected from a farm environment where the soil had never been developed or polluted. Soil interventions continued for eight weeks and all three mouse groups received aerosolized OVA/PBS challenge again on day 84. Mice were killed on day 85 and samples were collected for subsequent experiments.

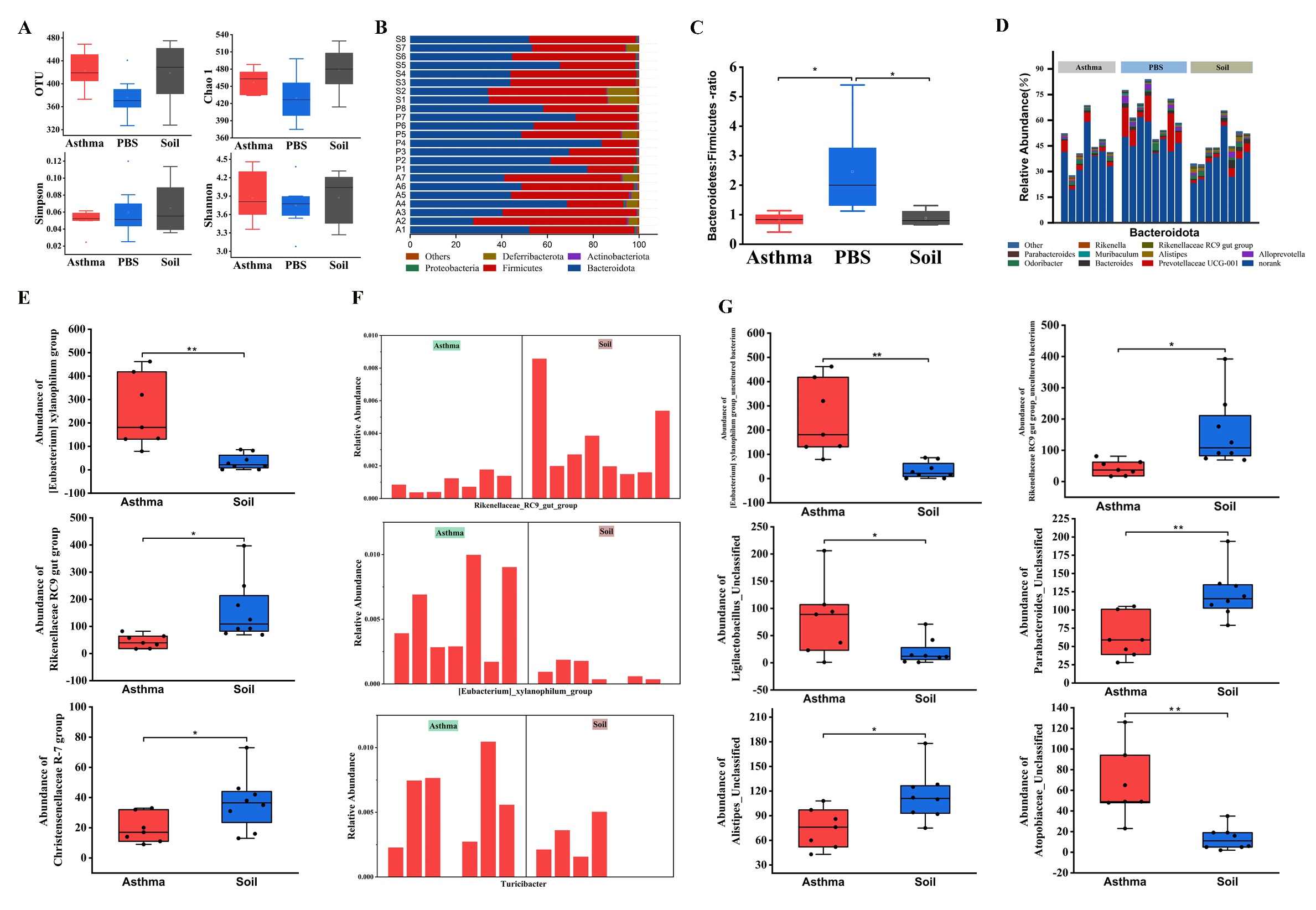
**Figure E3**

**Figure E3. Soil intake affected the structure and composition of intestinal microorganisms in asthmatic mice.** A. Richness and diversity comparison of the three experimental groups of mice. B. Percentage of gut bacterial community abundance on the phylum level after soil intervention. C. Soil intake did not influence the Bacteroidete/Firmicutes ratio. D. Effects of soil intake on fecal gut microbiota of the phylum Bacteroidota. E. Differential bacterial genera between the Asthma and Soil groups. F. Abundance of differentially abundant genera identified using LEfSe in the Asthma and Soil groups. G. Species abundance in the Asthma and Soil groups. Two-tailed least significant difference test, one-way ANOVA and non-parametric Mann–Whitney U test, Kruskal–Wallis test, and a post-hoc test with Bonferroni adjustment were performed. **P <* 0.05, ***P <* 0.01. Feces samples: PBS, n = 8; Asthma, n = 15.

**Figure E4**

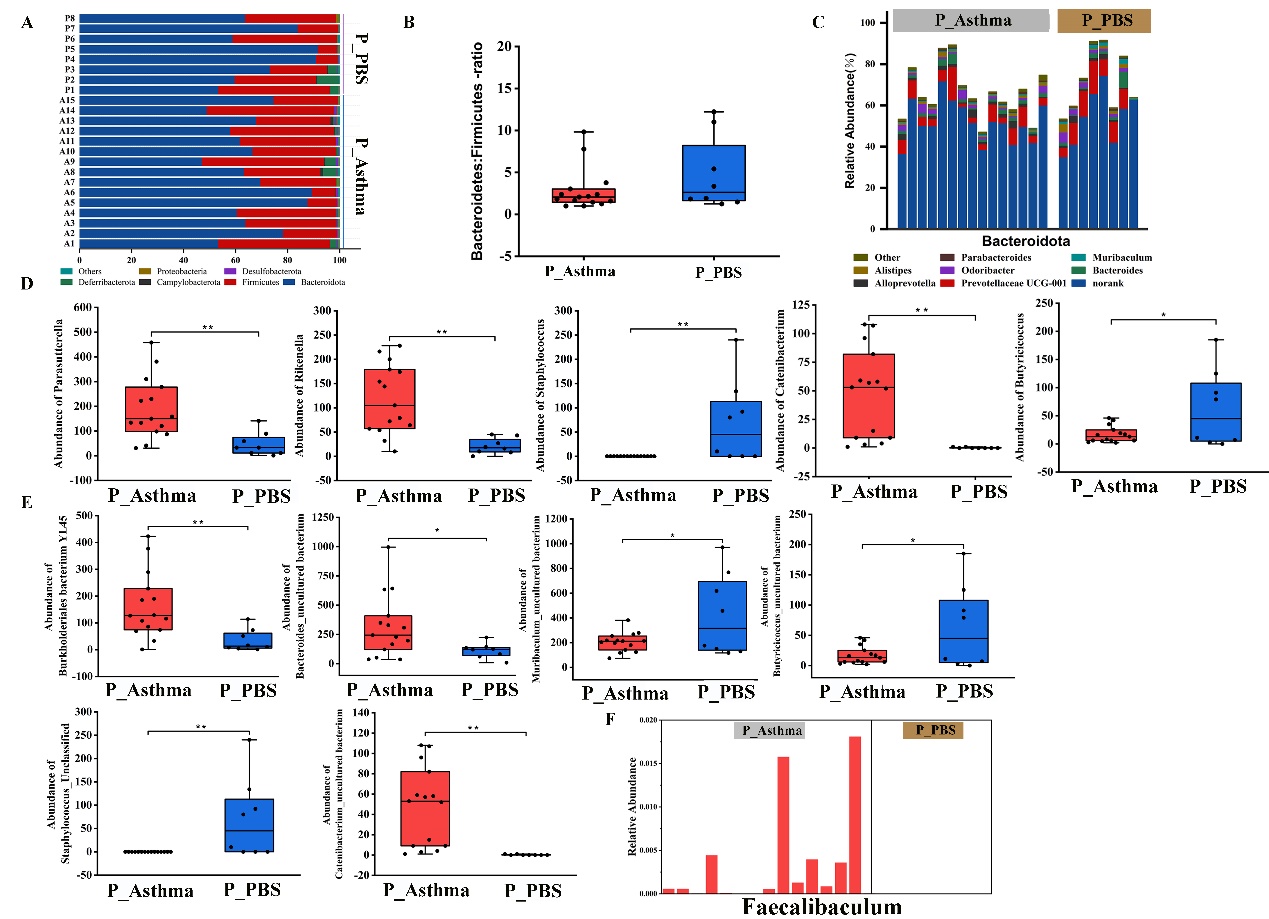

**Figure E4.** **Effects of asthma on intestinal microorganisms at the phylum, genus, and species level.** A. The percentage of gut bacterial community abundance on the phylum level. B. No effect of asthma on the Bacteroidete/Firmicutes ratio was observed. C. The relative abundance of the gut bacterial community on phylum Bacteroidetes. D. Differential bacterial genera between the two groups. E. Differential bacterial species. F. Relative abundance of genus *Faecalibaculum* for the most differentially abundant genera identified using LFfSe. Two-tailed least significant difference test. **P <* 0.05, ***P <* 0.01. Feces samples: P_PBS, n = 8; P_Asthma, n = 15.

**Figure E5**

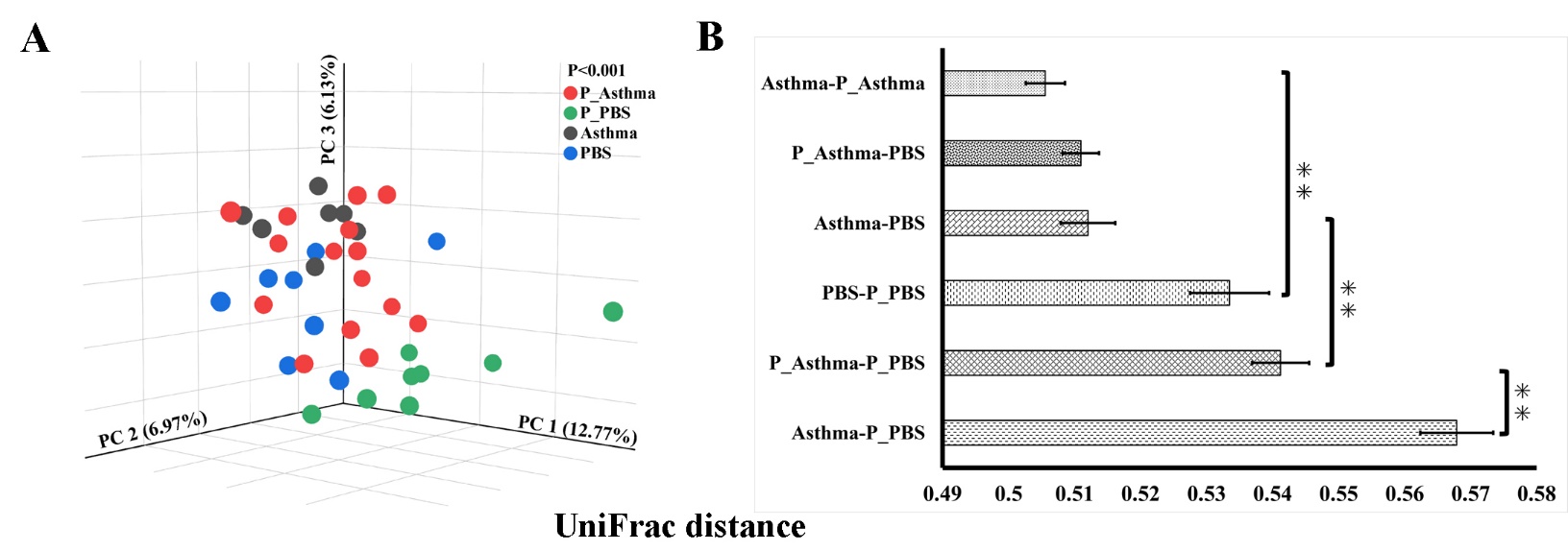

**Figure E5. Beta diversity of P_PBS, PBS, Asthma and P_Asthma groups.** A. PCoA of unweighted UniFrac distance. B. Each bar represents the mean ± SEM using unweighted UniFrac distance. Non-parametric Kruskal-Wallis test with Bonferroni post hoc. **P <* 0.05, ***P <* 0.01. n = 7-15.

**Figure E6**

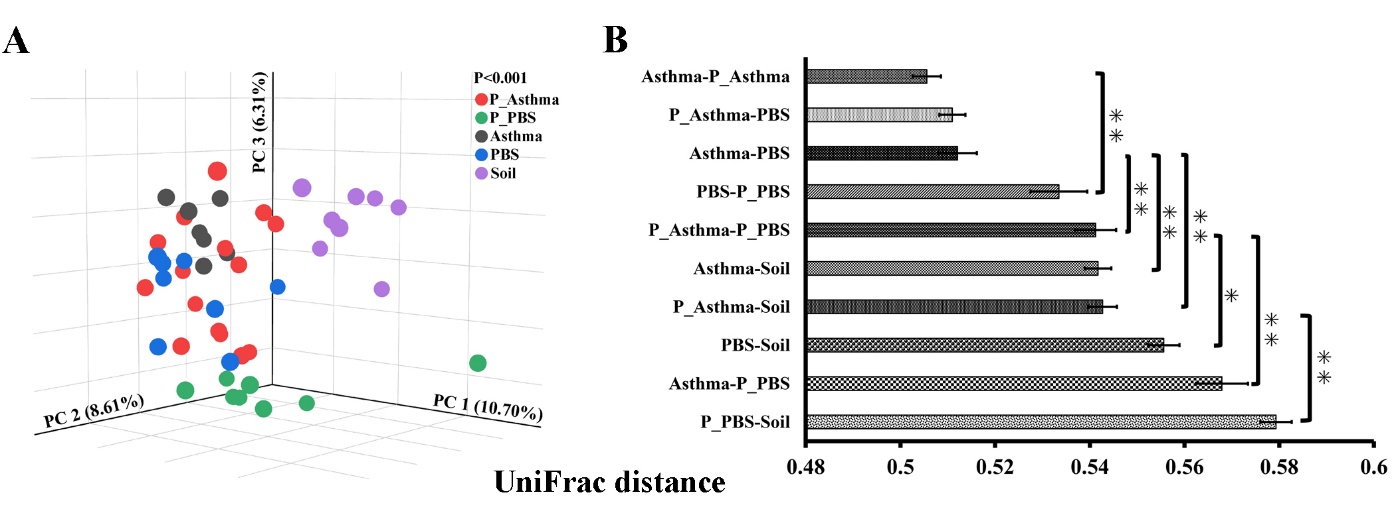

**Figure E6. Beta diversity of P_PBS, PBS, Asthma, P_Asthma and Soil groups.** A. PCoA of unweighted UniFrac distance. B. Each bar represents the mean ± SEM using the non-parametric Kruskal-Wallis test with Bonferroni post hoc. **P <* 0.05, ***P <* 0.01. n = 7-15.

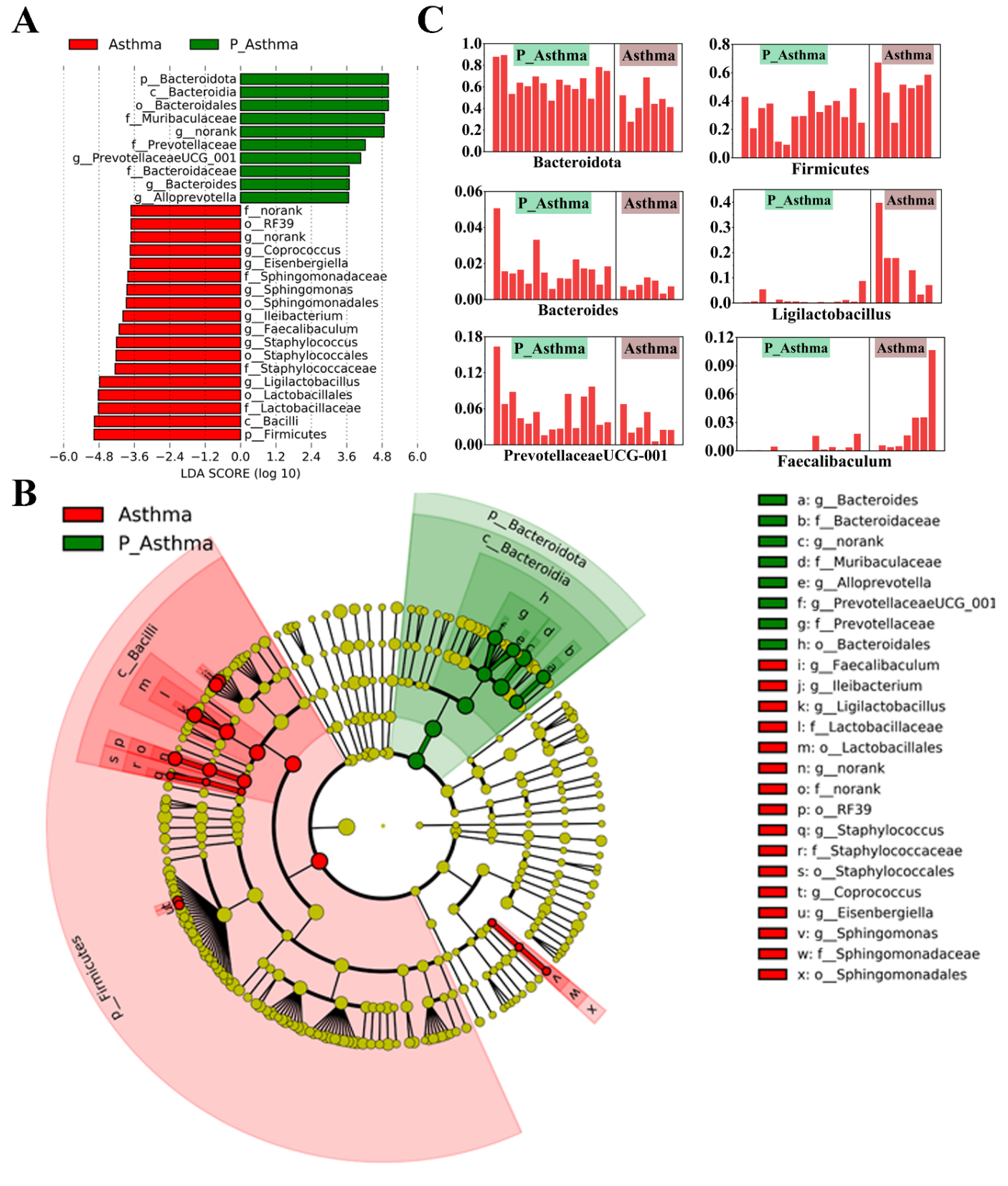
 **Figure E7**

**Figure E7. LEfSe analysis results on P_Asthma and Asthma.** A. Taxa enriched in P_Asthma are shown in green with positive LDA score and Asthma in red with negative LDA score (>3.5). B. Taxonomic cladogram from LEfSe. C. Differentially abundant taxa identified by LFfSe. n = 8-15.

**Figure E8**

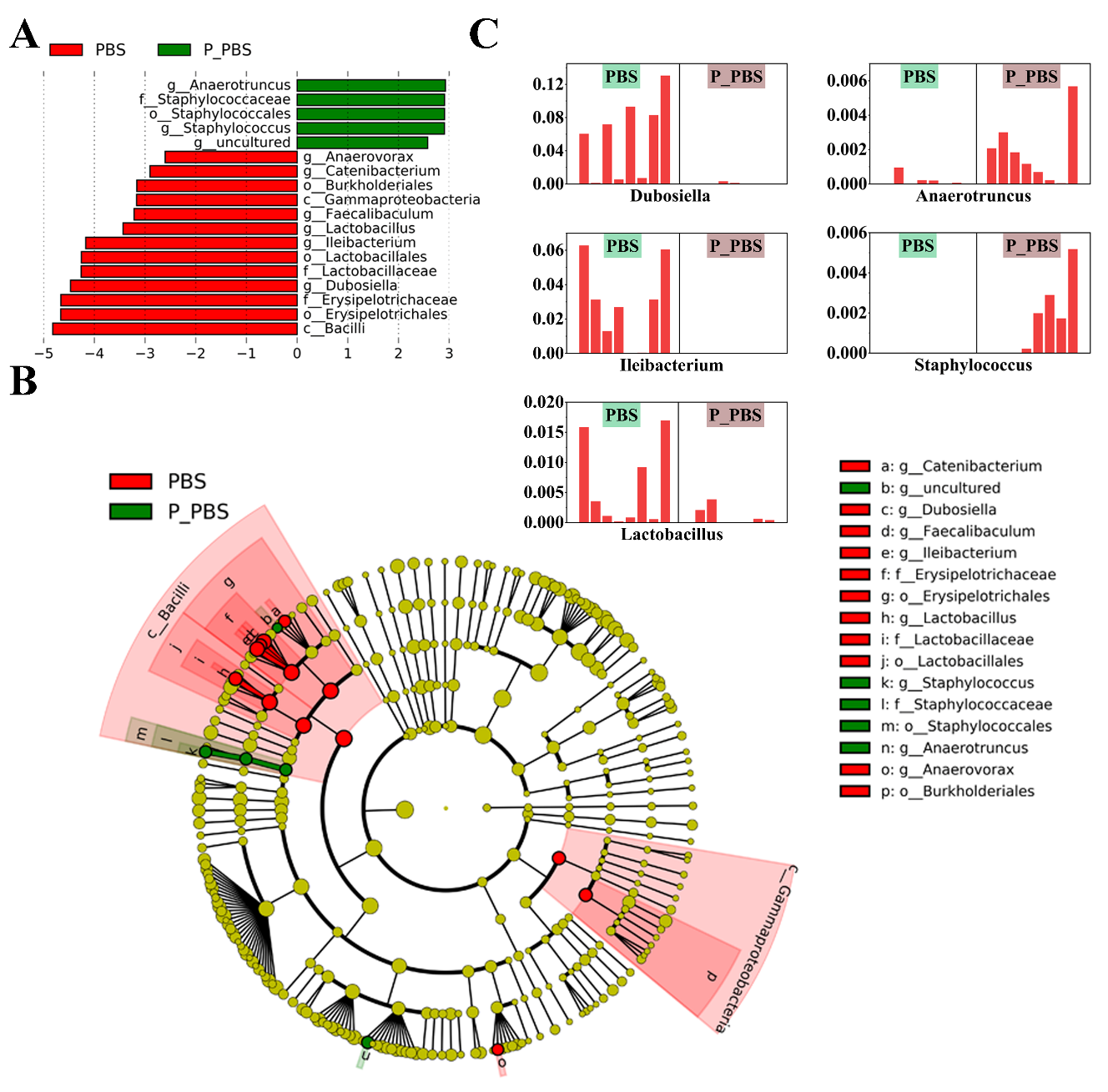

**Figure E8. LEfSe analysis results on P_PBS and PBS.** A. Taxa enriched in P_PBS are shown in green with positive LDA score and PBS in red with negative LDA score (>2.5). B. Taxonomic cladogram from LEfSe. C. Differentially abundant taxa identified by LFfSe. n = 8.

**Supplementary tables**

**Table E1**

| **Table E1. Mice and treatments** | | | | | | | | | |
| --- | --- | --- | --- | --- | --- | --- | --- | --- | --- |
| **SPF Animal Room** | | | | | | | | | |
| **SampleID** | **Treatment** | **Group** | **Reads** | **Diet** | **Water** | **Gender** | **Age** | **Age of Sample** | **Samples** |
| C11 | OVA+alum | P_Asthma | 57629 | Standard Diet | Sterilized Water | Male | 8-week（d0） | d28 | Feces |
| C12 | OVA+alum | P_Asthma | 51403 | Standard Diet | Sterilized Water | Male | 8-week（d0） | d28 | Feces |
| C13 | OVA+alum | P_Asthma | 59069 | Standard Diet | Sterilized Water | Male | 8-week（d0） | d28 | Feces |
| C14 | OVA+alum | P_Asthma | 47225 | Standard Diet | Sterilized Water | Male | 8-week（d0） | d28 | Feces |
| C15 | OVA+alum | P_Asthma | 50141 | Standard Diet | Sterilized Water | Male | 8-week（d0） | d28 | Feces |
| C16 | OVA+alum | P_Asthma | 49887 | Standard Diet | Sterilized Water | Male | 8-week（d0） | d28 | Feces |
| C17 | OVA+alum | P_Asthma | 46693 | Standard Diet | Sterilized Water | Male | 8-week（d0） | d28 | Feces |
| T11 | OVA+alum | P_Asthma | 66911 | Standard Diet | Sterilized Water | Male | 8-week（d0） | d28 | Feces |
| T12 | OVA+alum | P_Asthma | 67327 | Standard Diet | Sterilized Water | Male | 8-week（d0） | d28 | Feces |
| T13 | OVA+alum | P_Asthma | 54828 | Standard Diet | Sterilized Water | Male | 8-week（d0） | d28 | Feces |
| T14 | OVA+alum | P_Asthma | 68361 | Standard Diet | Sterilized Water | Male | 8-week（d0） | d28 | Feces |
| T15 | OVA+alum | P_Asthma | 50115 | Standard Diet | Sterilized Water | Male | 8-week（d0） | d28 | Feces |
| T16 | OVA+alum | P_Asthma | 63981 | Standard Diet | Sterilized Water | Male | 8-week（d0） | d28 | Feces |
| T17 | OVA+alum | P_Asthma | 61947 | Standard Diet | Sterilized Water | Male | 8-week（d0） | d28 | Feces |
| T18 | OVA+alum | P_Asthma | 46362 | Standard Diet | Sterilized Water | Male | 8-week（d0） | d28 | Feces |
| P11 | PBS | P_PBS | 48774 | Standard Diet | Sterilized Water | Male | 8-week（d0） | d28 | Feces |
| P12 | PBS | P_PBS | 47875 | Standard Diet | Sterilized Water | Male | 8-week（d0） | d28 | Feces |
| P13 | PBS | P_PBS | 53090 | Standard Diet | Sterilized Water | Male | 8-week（d0） | d28 | Feces |
| P14 | PBS | P_PBS | 48389 | Standard Diet | Sterilized Water | Male | 8-week（d0） | d28 | Feces |
| P15 | PBS | P_PBS | 56980 | Standard Diet | Sterilized Water | Male | 8-week（d0） | d28 | Feces |
| P16 | PBS | P_PBS | 63321 | Standard Diet | Sterilized Water | Male | 8-week（d0） | d28 | Feces |
| P17 | PBS | P_PBS | 57921 | Standard Diet | Sterilized Water | Male | 8-week（d0） | d28 | Feces |
| P18 | PBS | P_PBS | 55300 | Standard Diet | Sterilized Water | Male | 8-week（d0） | d28 | Feces |
| **Transfer all groups of mice to the General Animal Room** | | | | | | | | | |
| C1 | —— | Asthma | 63168 | Standard Diet | Sterilized Water | Male | 12-week（d28） | d85 | BALF, Lung, Feces, Serum |
| C2 | —— | Asthma | 52489 | Standard Diet | Sterilized Water | Male | 12-week（d28） | d85 | BALF, Lung, Feces, Serum |
| C3 | —— | Asthma | 56514 | Standard Diet | Sterilized Water | Male | 12-week（d28） | d85 | BALF, Lung, Feces, Serum |
| C4 | —— | Asthma | 68047 | Standard Diet | Sterilized Water | Male | 12-week（d28） | d85 | BALF, Lung, Feces, Serum |
| C5 | —— | Asthma | 59107 | Standard Diet | Sterilized Water | Male | 12-week（d28） | d85 | BALF, Lung, Feces, Serum |
| C6 | —— | Asthma | 49614 | Standard Diet | Sterilized Water | Male | 12-week（d28） | d85 | BALF, Lung, Feces, Serum |
| C7 | —— | Asthma | 58501 | Standard Diet | Sterilized Water | Male | 12-week（d28） | d85 | BALF, Lung, Feces, Serum |
| P1 | —— | PBS | 58515 | Standard Diet | Sterilized Water | Male | 12-week（d28） | d85 | BALF, Lung, Feces, Serum |
| P2 | —— | PBS | 59450 | Standard Diet | Sterilized Water | Male | 12-week（d28） | d85 | BALF, Lung, Feces, Serum |
| P3 | —— | PBS | 53527 | Standard Diet | Sterilized Water | Male | 12-week（d28） | d85 | BALF, Lung, Feces, Serum |
| P4 | —— | PBS | 55934 | Standard Diet | Sterilized Water | Male | 12-week（d28） | d85 | BALF, Lung, Feces, Serum |
| P5 | —— | PBS | 62431 | Standard Diet | Sterilized Water | Male | 12-week（d28） | d85 | BALF, Lung, Feces, Serum |
| P6 | —— | PBS | 54297 | Standard Diet | Sterilized Water | Male | 12-week（d28） | d85 | BALF, Lung, Feces, Serum |
| P7 | —— | PBS | 62145 | Standard Diet | Sterilized Water | Male | 12-week（d28） | d85 | BALF, Lung, Feces, Serum |
| P8 | —— | PBS | 60378 | Standard Diet | Sterilized Water | Male | 12-week（d28） | d85 | BALF, Lung, Feces, Serum |
| T1 | —— | Soil | 58266 | Standard Diet Containing 5% Sterilized Soil | Sterilized Water | Male | 12-week（d28） | d85 | BALF, Lung, Feces, Serum |
| T2 | —— | Soil | 58855 | Standard Diet Containing 5% Sterilized Soil | Sterilized Water | Male | 12-week（d28） | d85 | BALF, Lung, Feces, Serum |
| T3 | —— | Soil | 58807 | Standard Diet Containing 5% Sterilized Soil | Sterilized Water | Male | 12-week（d28） | d85 | BALF, Lung, Feces, Serum |
| T4 | —— | Soil | 58858 | Standard Diet Containing 5% Sterilized Soil | Sterilized Water | Male | 12-week（d28） | d85 | BALF, Lung, Feces, Serum |
| T5 | —— | Soil | 60607 | Standard Diet Containing 5% Sterilized Soil | Sterilized Water | Male | 12-week（d28） | d85 | BALF, Lung, Feces, Serum |
| T6 | —— | Soil | 57876 | Standard Diet Containing 5% Sterilized Soil | Sterilized Water | Male | 12-week（d28） | d85 | BALF, Lung, Feces, Serum |
| T7 | —— | Soil | 58642 | Standard Diet Containing 5% Sterilized Soil | Sterilized Water | Male | 12-week（d28） | d85 | BALF, Lung, Feces, Serum |
| T8 | —— | Soil | 53381 | Standard Diet Containing 5% Sterilized Soil | Sterilized Water | Male | 12-week（d28） | d85 | BALF, Lung, Feces, Serum |

**Table E2**

| **Table E2 (A). P values of multiple comparisons of unweighted UniFrac distance between bacterial fecal communities of mice fed different diets.** | | | |
| --- | --- | --- | --- |
|  | PBS-Soil | Asthma-Soil | Asthma-PBS |
| PBS-Soil | 0 | 0.057 | 0.000 |
| Asthma-Soil |  | 0 | 0.000 |
| Asthma-PBS |  |  | 0 |
| **Table E2 (B). P values of multiple comparisons of Bray-Curtis between bacterial fecal communities of mice fed different diets.** | | | |
|  | PBS-Soil | Asthma-Soil | Asthma-PBS |
| PBS-Soil | 0 | 0.198 | 0.000 |
| Asthma-Soil |  | 0 | 0.027 |
| Asthma-PBS |  |  | 0 |
| P values were based on one-way ANOVA. | | | |

**Table E3**

| **Table E3. Main phyla in each group after 8 weeks of soil intervention.** | | | | | | |
| --- | --- | --- | --- | --- | --- | --- |
| **Phyla** | **Mean** | | | **Significance** | | |
|  | **PBS** | **Asthma** | **Soil** | **Soil-PBS** | **Soil-Asthma** | **Asthma-PBS** |
| Bacteroidota | 0.6575 | 0.4615 | 0.4651 | **※※** | —— | **※※** |
| Firmicutes | 0.3240 | 0.4971 | 0.4848 | **※※** | —— | **※※** |
| Deferribacterota | 0.0086 | 0.0272 | 0.0387 | —— | —— | —— |
| Actinobacteriota | 0.0044 | 0.0045 | 0.0045 | —— | **※※** | **※※** |
| Proteobacteria | 0.0029 | 0.0055 | 0.0031 | —— | —— | —— |
| Campylobacterota | 0.0011 | 0.0022 | 0.0016 | —— | —— | —— |
| Desulfobacterota | 0.0003 | 0.0006 | 0.0016 | —— | —— | —— |
| Cyanobacteria | 0.0009 | 0.0005 | 0.0001 | —— | —— | —— |
| ^※^*P* < 0.05, ^※※^*P* < 0.01. P values were based on one-way ANOVA and non-parametric Kruskal-Wallis test. | | | | | | |

**Table E4**

| **Table E4. Top 20 most abundant genera in the three groups after soil intervention.** | | | | | | | |
| --- | --- | --- | --- | --- | --- | --- | --- |
| **Genus** | **Mean** | | | **P-value** | | | **Statistical Methods** |
|  | **PBS** | **Asthma** | **Soil** | **Soil-PBS** | **Soil-Asthma** | **Asthma-PBS** |  |
| Muribaculaceae_norank | 0.4947 | 0.3804 | 0.3588 | **※** | —— | **※** | ANOVA |
| Ligilactobacillus | 0.0429 | 0.1412 | 0.0355 | —— | **※** | **※** | ANOVA |
| Dubosiella | 0.0565 | 0.0397 | 0.0754 | —— | —— | —— | KW |
| Prevotellaceae UCG-001 | 0.0939 | 0.0322 | 0.0296 | —— | —— | —— | KW |
| Lachnospiraceae NK4A136 group | 0.0450 | 0.0553 | 0.0507 | —— | —— | —— | ANOVA |
| Ileibacterium | 0.0282 | 0.0434 | 0.0745 | —— | —— | —— | KW |
| Clostridia UCG-014_norank | 0.0300 | 0.0459 | 0.0468 | —— | —— | —— | KW |
| Mucispirillum | 0.0086 | 0.0272 | 0.0387 | —— | —— | —— | KW |
| Lachnospiraceae_uncultured | 0.0147 | 0.0199 | 0.0328 | —— | —— | —— | KW |
| RF39_norank | 0.0197 | 0.0219 | 0.0143 | —— | —— | —— | ANOVA |
| Bacteroides | 0.0154 | 0.0077 | 0.0188 | —— | —— | —— | KW |
| Allobaculum | 0.0000 | 0.0000 | 0.0407 | **※※** | **※※** | —— | KW |
| Oscillospiraceae_uncultured | 0.0124 | 0.0114 | 0.0181 | —— | —— | —— | ANOVA |
| Faecalibaculum | 0.0030 | 0.0297 | 0.0100 | —— | —— | **※** | KW |
| Odoribacter | 0.0108 | 0.0143 | 0.0150 | —— | —— | —— | ANOVA |
| Alloprevotella | 0.0185 | 0.0072 | 0.0100 | —— | —— | —— | KW |
| Colidextribacter | 0.0079 | 0.0101 | 0.0173 | —— | —— | —— | KW |
| Lactobacillus | 0.0060 | 0.0193 | 0.0038 | —— | —— | —— | KW |
| Alistipes | 0.0057 | 0.0065 | 0.0148 | **※※** | **※** | —— | KW |
| Limosilactobacillus | 0.0145 | 0.0017 | 0.0017 | —— | —— | —— | KW |
| ^※^*P* < 0.05, ^※※^*P* < 0.01. ANOVA was used when the variances were homogeneous, and a non-parametric test (Kruskal-Wallis) for non-homogeneous variances. | | | | | | | |

**Table E5**

| **Table E5. Differential species with sum of abundance greater than 500 in the Asthma and Soil groups after soil intervention.** | | | | |
| --- | --- | --- | --- | --- |
| **Species** | **Mean** | | **Total** | **P-value** |
|  | **Asthma** | **Soil** |  |  |
| Alistipes_uncultured bacterium | 225.000 | 569.375 | 6130.000 | 0.018 |
| Lachnospiraceae UCG-001_uncultured bacterium | 19.429 | 329.875 | 2775.000 | 0.030 |
| Turicibacter_uncultured bacterium | 238.571 | 71.250 | 2240.000 | 0.030 |
| [Eubacterium] xylanophilum group_uncultured bacterium | 246.429 | 34.000 | 1997.000 | **0.002** |
| Rikenellaceae RC9 gut group_uncultured bacterium | 43.286 | 158.000 | 1567.000 | 0.020 |
| Alistipes_Unclassified | 74.571 | 114.250 | 1436.000 | 0.017 |
| Parabacteroides_Unclassified | 62.429 | 122.250 | 1415.000 | **0.003** |
| UCG-009_uncultured bacterium | 76.000 | 30.500 | 776.000 | **0.006** |
| Ligilactobacillus_Unclassified | 79.571 | 20.500 | 721.000 | 0.039 |
| Atopobiaceae_Unclassified | 64.857 | 13.375 | 561.000 | **0.001** |
| P values were based on a two-tailed least significant difference test. | | | | |

**Table E6**

| **Table E6. Comparison of relative mRNA expression of cytokines in lung tissue of three groups of mice after soil intake.** | | | |
| --- | --- | --- | --- |
| **Cytokines** | **Group** | | |
|  | **PBS** | **Asthma** | **Soil** |
| IL-4 | 1.27±0.86 | 8.55±5.59** | 3.44±1.11* |
| IL-5 | 1.05±0.36 | 1.13±0.19 | 0.99±0.48 |
| IL-10 | 1.08±1.16 | 1.17±0.41 | 4.86±3.55 |
| IL-17A | 1.35±1.04 | 3.71±1.31* | 3.39±1.03* |
| IL-17F | 1.02±0.23 | 1.3±0.23 | 1.2±0.38 |
| IFN-γ | 1.89±1.26 | 1.3±0.41 | 5.93±3.63▲ |
| Th1/Th2 | 1.66±1.26 | 0.72±0.12 | 2.18±2.23 |
| Treg/Th17 | 2.17±2.31 | 0.91±0.33 | 3.78±2.33 |
| Treg/Th2 | 2.42±2.67 | 1.05±0.36 | 6.82±6 |
| P values were based on non-parametric test (Kruskal-Wallis).  * Compared with PBS group. ^*^*P* < 0.05, ^**^*P* < 0.01. ▲ Compared with Asthma group. ^▲^*P* < 0.05, ^▲▲^*P* < 0.01. | | | |

**Table E7**

| **Table E7. Major phyla of P_Asthma and P_PBS groups.** | | | |
| --- | --- | --- | --- |
| **Phyla** | **Mean** | | **P-value** |
|  | **P_Asthma** | **P_PBS** |  |
| Bacteroidota | 0.6606 | 0.7196 | 0.329 |
| Firmicutes | 0.3162 | 0.2530 | 0.262 |
| Deferribacterota | 0.0103 | 0.0189 | 0.376 |
| Actinobacteriota | 0.0037 | 0.0035 | 0.912 |
| Proteobacteria | 0.0046 | 0.0014 | 0.006 |
| Campylobacterota | 0.0023 | 0.0015 | 0.552 |
| Desulfobacterota | 0.0011 | 0.0014 | 0.699 |
| Cyanobacteria | 0.0007 | 0.0006 | 0.784 |
| P values were based on a two-tailed least significant difference test. | | | |

**Table E8**

| **Table E8. Top 20 most abundant genera in P_Asthma and P_PBS groups.** | | | |
| --- | --- | --- | --- |
| **Genus** | **Mean** | | **P-value** |
|  | **P_PBS** | **P_Asthma** |  |
| Muribaculaceae_norank | 0.5208 | 0.5438 | 0.651 |
| Prevotellaceae UCG-001 | 0.0587 | 0.0888 | 0.123 |
| Lachnospiraceae NK4A136 group | 0.0610 | 0.0714 | 0.670 |
| Clostridia UCG-014_norank | 0.0291 | 0.0246 | 0.555 |
| Dubosiella | 0.0379 | 0.0006 | **0.007** |
| Lachnospiraceae_uncultured | 0.0219 | 0.0235 | 0.791 |
| Ileibacterium | 0.0341 | 0.0000 | 0.179 |
| Odoribacter | 0.0182 | 0.0200 | 0.775 |
| Bacteroides | 0.0177 | 0.0204 | 0.724 |
| Ligilactobacillus | 0.0139 | 0.0275 | 0.433 |
| Oscillospiraceae_uncultured | 0.0159 | 0.0181 | 0.730 |
| Alloprevotella | 0.0167 | 0.0134 | 0.489 |
| Mucispirillum | 0.0103 | 0.0189 | 0.376 |
| RF39_norank | 0.0120 | 0.0113 | 0.799 |
| Colidextribacter | 0.0108 | 0.0127 | 0.629 |
| Alistipes | 0.0096 | 0.0130 | 0.369 |
| Lachnoclostridium | 0.0093 | 0.0071 | 0.704 |
| Muribaculum | 0.0053 | 0.0105 | **0.025** |
| [Eubacterium] xylanophilum group | 0.0060 | 0.0079 | 0.650 |
| Parabacteroides | 0.0050 | 0.0048 | 0.873 |
| P values were based on a two-tailed least significant difference test. | | | |

**Table E9**

| **Table E9. Differential species with sum of abundance greater than 500 in the P_Asthma and P_PBS groups.** | | | | |
| --- | --- | --- | --- | --- |
| **Species** | **Mean** | | **Total** | **P-value** |
|  | **P_Asthma** | **P_PBS** |  |  |
| Dubosiella_uncultured bacterium | 1753.733 | 11.250 | 26396.000 | **0.007** |
| Muribaculum_uncultured bacterium | 205.467 | 423.250 | 6468.000 | 0.022 |
| Bacteroides_uncultured bacterium | 316.400 | 112.000 | 5642.000 | 0.046 |
| Burkholderiales bacterium YL45 | 162.333 | 34.750 | 2713.000 | **0.010** |
| Rikenella_Unclassified | 55.533 | 0.000 | 833.000 | **0.008** |
| Butyricicoccus_uncultured bacterium | 16.600 | 62.625 | 750.000 | 0.019 |
| Catenibacterium_uncultured bacterium | 47.533 | 0.250 | 715.000 | **0.003** |
| Staphylococcus_Unclassified | 0.000 | 69.500 | 556.000 | **0.004** |
| UCG-010_uncultured bacterium | 28.267 | 10.125 | 505.000 | 0.028 |
| P values were based on a two-tailed least significant difference test. | | | | |
